# Scaled-expansion of the membrane associated cytoskeleton requires conserved kinesin adaptors

**DOI:** 10.1101/2022.05.09.491207

**Authors:** Oliver Glomb, Grace Swaim, Pablo Munoz LLancao, Christopher Lovejoy, Sabyasachi Sutradhar, Junhyun Park, Youjun Wu, Marc Hammarlund, Jonathon Howard, Shawn M. Ferguson, Shaul Yogev

## Abstract

A periodic lattice of actin rings and spectrin tetramers scaffolds the axonal membrane. How spectrin is delivered to this structure to scale its size to that of the growing axon is unknown. We found that endogenous spectrin, visualized with singe axon resolution *in vivo*, is delivered to hotspots in the lattice that support its expansion at rates set by axon stretch-growth. Unlike other cytoskeletal proteins, whose apparent slow movement consists of intermittent bouts of fast movements, spectrin moves slowly and processively. We identified a pair of coiled coil proteins that mediate this slow movement and the expansion of the lattice by linking spectrin to kinesin-1. Thus, processive slow transport and local lattice incorporation support scaled cytoskeletal expansion during axon stretch-growth.

**One-Sentence Summary:** Kinesin adaptors control spectrin transport and expansion of the membrane periodic skeleton.

## Introduction

A periodic lattice of actin rings connected by spectrin tetramers forms a Membrane associated Periodic Skeleton (MPS) that underlies the axonal membrane^1, 2^. During early axon outgrowth in culture, the MPS extends distally from the cell body^3^, likely by addition of subunits at its distal end. *In vivo,* early axon outgrowth is followed by a process termed interstitial elongation or stretch-growth, which accounts for most of the mature axon length^4^. During this stage, the axon elongates without a growth cone at a rate that matches organismal growth. To scale to the size of the elongating axon during stretch-growth, the MPS must expand, presumably, by incorporating newly delivered spectrin tetramers and other cytoskeletal proteins. How this expansion happens is unknown.

Radiolabeling newly synthesized proteins during axon stretch growth revealed an apparent slow progression of spectrin (0.7-8 mm/day, with a maximal rate of 50 mm/day) from the cell body to the axon^5–7^. Although the mechanisms underlying this progression are unknown, it suggests that slow delivery of spectrin supports MPS expansion. Directly visualizing slow-moving cytoskeletal and soluble proteins in the axon with fluorescence-based approaches is challenging due to their slow movement and ubiquitous presence^8^. Interestingly, fluorescent labelling of slow transport cargo such as neurofilaments, microtubules, actin, dynein, synapsin or clathrin does not show processive slow movements, but rather intermittent fast transport or biased diffusion underlined by fast transport^8–14^. Hence, current models suggest that the observed slow vectorial movement in radiolabeling studies in fact represents intermittent fast transport^8, 12^. The basis for such intermittent fast transport remains to be established for most slow axonal transport cargos. Furthermore, as many slow transport cargos have never been visualized directly or investigated mechanistically, it is also possible that additional mechanisms exist.

## Results

To investigate the dynamic processes that drive MPS (Membrane associated Periodic Skeleton) expansion *in vivo*, we built a probe for temporally controlled labeling of endogenous, newly synthesized spectrin, with single axon resolution in *C. elegans* (Fig. 1A, B). We engineered an FRT-flanked termination sequence, followed by a 7xspGFP11 tag into the C-terminus of *spc-1* – the sole worm α-spectrin – to generate *spc-1^Flip-On^::GFP.* We drove FLP expression with a heat shock promoter for temporal control of labeling and visualized SPC-1^Flip-On^::GFP specifically in the DA9 motor neuron by expressing spGFP1-10 with the *mig-13* promoter (Fig. 1B). A constitutively expressed SPC-1::GFP was shown to be functional in worms^15^ and the MPS in DA9 was shown to have stereotypical periodicity^16^. We confirmed that SPC-1::GFP integrates into the MPS lattice using FRAP experiments, which in our hands were a more robust readout for MPS integrity compared to measuring periodicity. As in hippocampal neurons^3^, SPC-1::GFP showed little recovery in FRAP, but recovered rapidly upon single cell degradation (*scKD*) of *unc-70/β-spectrin* (Fig. S1).

**Figure 1:**
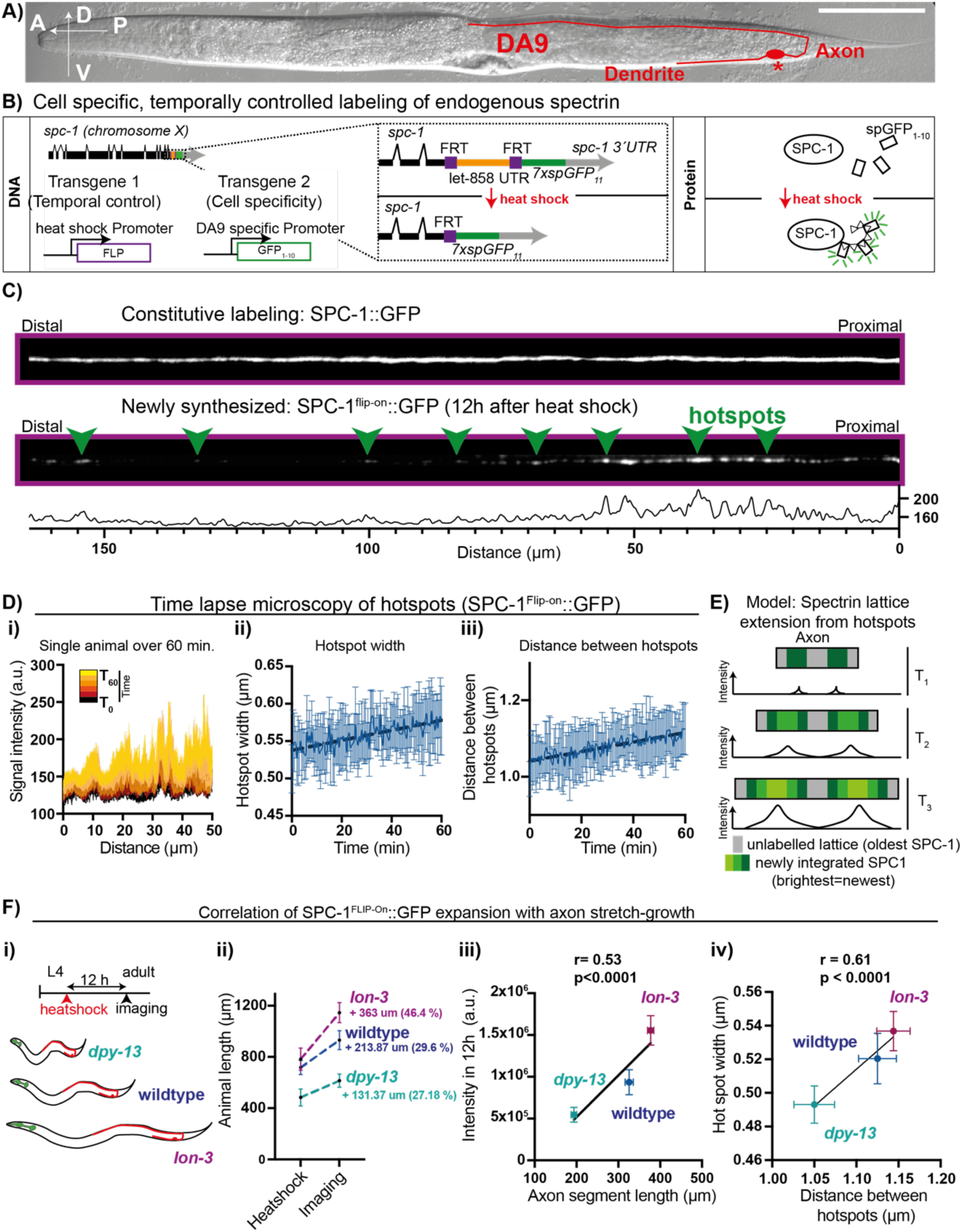
MPS lattice expansion from hotspots during axon stretch-growth. A. Schematic of the DA9 neuron (red, * marks the cell body) in an L4 animal. Scale bar: 100 μm. B. Labelling strategy for single cell pulse labeling of endogenous α-spectrin (SPC-1^Flip-On^::GFP). C. Comparison of constitutive (SPC-1::GFP) and pulse expression of spectrin (SPC-1^Flip-On^::GFP) at 12 hrs after heatshock induction. Arrowheads point to hotspots where newly synthesized spectrin accumulates. Scale:10 μm. D. Time lapse microscopy of hotspot dynamics at 8h after heat shock (n=9). i) Intensity profile of SPC-1^Flip-on^::GFP in a single animal shows that signal increases over time in the same locations. Time is color-coded from dark (T=0) to light (T=60). ii) Width at half maximum of hot spots and iii) the distance between hotspot center intensity increases over one hour. E. Proposed model for MPS lattice expansion during axon stretch-growth by expansion of spectrin from hotspots. F. Spectrin expansion correlates with axon stretch-growth. SPC-1 signal was measured 12h after heat shock in wildtype and body-size mutants *lon-3(e2175)* and *dpy-13(e184)* animals (i). Animals were heat shocked in late L4 and imaged as young adults. ii) Animal length was measured at the time of heatshock or at imaging 12h after heat shock. Growth increment in μm or as % are indicated. iii) Levels of SPC-1^Flip-On^::GFP produced in 12 hrs correlate with increase in axon length (measured between the turn of the commissure and the cell body of SDQL) in body-size mutants. iv) Hotspot width and distance between hotspots increase in correlation to each other over 12hrs across the three genotypes. Pearson r and p-value as indicated (iii+iv).

Surprisingly, despite the uniform distribution of the MPS and its growth from its distal end during early development, newly synthesized spectrin localized to discrete puncta at roughly stereotypic positions along the length of the axon (Fig 1F), suggesting that they might represent hot spots for spectrin incorporation and MPS lattice expansion at later stages. The puncta were mostly immobile and did not recover in FRAP experiments (Fig. S1C), confirming that they represent SPC-1 in the MPS lattice. To understand how the dynamics of these hotspots might support MPS expansion, we followed SPC-1^Flip-On^::GFP puncta at 8 hours post induction by live imaging. The intensity of individual puncta increased over one hour, consistent with their proposed function as hotspots for integrating new spectrin into the lattice (Fig. 1J). Hot spots grew in width and their intensity peaks moved apart from each other (Fig 1H), suggesting that the MPS expands from these sites (Fig 1E). These data suggest that local incorporation of spectrin at hotspots could underlie MPS expansion. What sets the rate of MPS expansion? The DA9 axon reaches its target during embryogenesis, after which its length scales with animal length in the absence of a growth cone, suggesting that it is undergoing stretch-growth. Consistent with the idea that stretch growth determines the rate of MPS expansion, we noticed that the rate of SPC-1^Flip-On^::GFP fluorescence increase (∼3.6 fold in 34 hrs) was roughly similar to the rate of animal anterior-posterior axis elongation (∼3.1 fold) (Fig. S2 A, C) and therefore the rate of DA9 stretch growth. SPC-1^Flip-On^::GFP fluorescence increased linearly, suggesting that at the observed timepoints, it reflects mostly the addition of newly synthesized molecules into the expanding MPS, rather than turnover of spectrin at steady-state (in which case we would expect the signal to plateau). To directly test if MPS expansion is driven by growth of the animal, we compared the intensity of newly synthesized SPC-1^Flip-On^::GFP during a 12 hour time window between wildtype and mutants that show reduced (*dpy-13(e184))* or increased (*lon-3(e2175))* body growth rates and hence axon stretch-growth rates (Fig. 1J_(ii)_). Importantly, DPY-13 and LON-3 are structural components of the worm’s cuticle^17, 18^, and therefore their effects on axon length are indirect and are driven by changes to the animal’s size. The levels of newly synthesized spectrin correlated well with the rate of animal growth across these genotypes, suggesting that the rate of MPS lattice expansion is driven by the axon’s stretch-growth (Fig. 1J_(iii)_).

To test if axon stretch growth promotes MPS expansion from hotspots, we compared hotspots dynamics in wildtype, *dpy-13* and *lon-3* animals (Fig. 1J, S2D). The total number of hotspots was mildly decreased in *dpy-13* mutants but did not change significantly in *lon-3* mutants compared to wildtype (Fig. S2E). Both puncta width and the distance between puncta peaks correlated with the change in axon length in body size mutants (Fig S2F,G). Importantly, hotspot width and the distance between hotspots correlated well across genotypes (Fig. 1J_(IV)_), strongly arguing that the process of axon stretch-growth, which depends on animal growth rates, drives expansion of the spectrin lattice from hotspots. Together, these results point to the existence of hot spots of spectrin integration into the lattice, which would locally sustain MPS expansion during stretch-growth of the axon, scaling the length of the cytoskeleton to that of the axon (Fig. 1J).

We next sought to understand the delivery of spectrin to hotspots. We tested how mutations in kinesin motors – several of which co-precipitate with spectrin from mouse brain^19^ – affect MPS distribution. Mutations in kinesin-3/KIF1A (*unc-104(e1265)*) did not affect the MPS (Fig S3A), whereas two mutant alleles of kinesin-1/KIF5 (*unc-116(e2310)* and *unc-116(rh24sb79)*) led to a strong reduction in distal SPC-1::GFP (Fig. 2A,B). Expression of *unc-116* cDNA in DA9 rescued MPS distribution in *unc-116* mutants and cell-specific degradation of UNC-116 phenocopied *unc-116* mutants, indicating that kinesin-1/KIF5 functions in DA9 to promote MPS formation (Fig 2B). *unc-116(e2310)* mutants show normal axon length, suggesting that kinesin-1/KIF5 functions to promote MPS formation independently of its role in early axon development.

**Figure 2:**
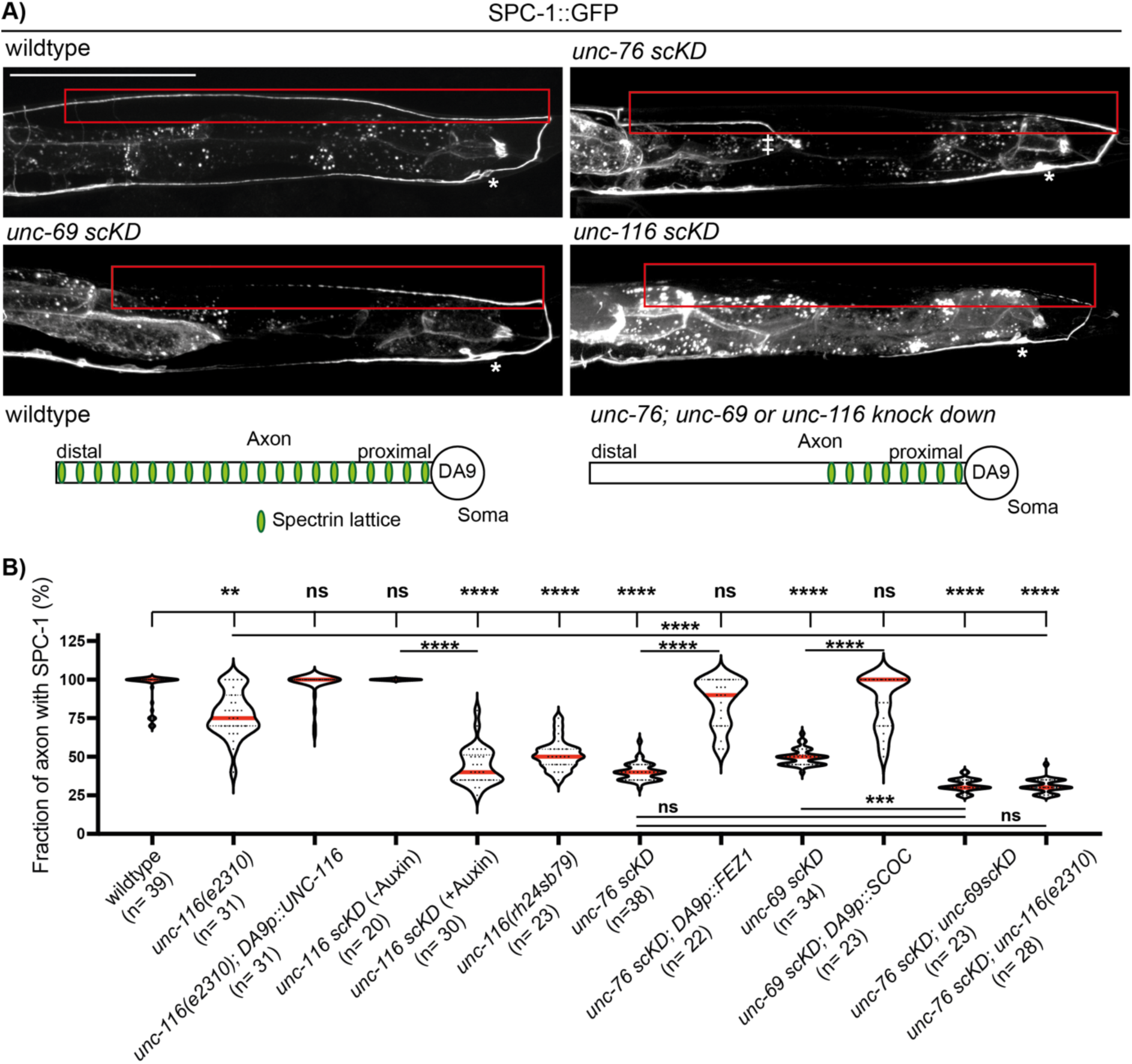
Kinesin-1/UNC-116, FEZ1/UNC-76 and SCOC/UNC-69 are required for spectrin lattice formation. A. Representative images of SPC-1::GFP in wildtype or single-cell knock down (*scKD*) of *unc-116*, *unc-76* or *unc-69* show strong loss of the spectrin signal in the axon, highlighted by boxes. * indicates the DA9 cell body. Scale: 100 μm. B. Quantification of the axonal fraction that contains SPC-1::GFP signal in indicated genotypes. Red line and faint black lines in violin indicate mean and 25^th^/75^th^ percentiles, respectively. ** p < 0.01; *** p < 0.001; **** p < 0.0001, based on a Kruskal-Wallis test followed by uncorrected Dunńs post-test for multiple comparisons.

To identify potential spectrin-kinesin adaptors, we tested mutations in the *C. elegans* orthologues of JIP1, JIP3 and kinesin light chains 1 and 2 but did not detect an effect on MPS distribution (Fig. S3A). Conversely, loss of the kinesin-1 binding protein UNC-76/FEZ1^20–23^ strongly reduced the distal MPS, similar to *unc-116* mutants (Fig. S4). This points to UNC-76/FEZ1 as being a potential kinesin-1 adaptor for spectrin, although we cannot entirely rule out a minor role for other adaptors, since the mutations tested were not null. We generated a cell specific degradation allele of *unc-76 (unc-76 scKD)* to circumvent the axon fasciculation defects observed in these mutants^23^ and saw a strong loss of the MPS (Fig 2A, B). The DA9 axon was ∼30% shorter in these animals than in wildtype (223 µm ±24 compared to 325 µm ±49, respectively), but axon guidance was not affected and no degeneration was observed at the L4 stage, suggesting that loss of the MPS is a direct outcome of reduced UNC-76 activity. Degradation of UNC-76 only slightly enhanced MPS loss in *unc-116* mutants, consistent with the two proteins functioning together and the two alleles not being complete nulls (Fig 2B).

To understand how UNC-76/FEZ1 functions, we tested whether any of its binding partners are required for MPS formation. We found that a single cell degradation allele of the poorly characterized short coiled-coil protein, UNC-69/SCOC^24^, phenocopied *unc-76* (Fig. 2A, B). Double degradation of both proteins only slightly enhanced the single degradation phenotypes, suggesting that both proteins function in the same pathway. Expression of the human homologs of UNC-76 and UNC-69, FEZ1 and SCOC, could rescue MPS defects in the respective mutants, indicating that the function of these adaptors is highly conserved (Fig 2B).

Tagging endogenous UNC-76 and UNC-69 with 10XspGFP11 or 7XspGFP11 revealed that they are localized uniformly throughout the DA9 axon (Fig. S5). We found that UNC-76 and UNC-69 depend on each other for their axonal localization, and confirmed that UNC-76 localization is *unc-116* dependent (Fig. S5)^24^. Together with previous reports of UNC-76/FEZ1 binding to kinesin^20^ and UNC-69 binding to UNC-76^24^, these results strongly argue that UNC-76, UNC-116 and UNC-69 function together to mediate MPS assembly. Consistent with this interpretation, other MPS components UNC-70/μ spectrin and UNC-44/Ankyrin were also lost from the distal axon upon loss of *unc-76* either in DA9 or in additional neurons (Fig. S4). Conversely, two other kinesin-1 cargo – mitochondria and RAB-11 vesicles – were not affected (Fig. S6), highlighting the specificity of this adaptor complex to spectrin.

Next, we used our conditional SPC-1^Flip-On^::GFP probe to visualize the slow transport of spectrin and how it may be affected in the mutants which led to MPS loss. Strikingly, SPC-1^Flip-On^::GFP movement was different than that described for other cytoskeletal proteins: instead of intermittent fast transport or biased diffusion, we observed slow processive runs (Fig 3A, B). The speed of these runs (∼0.009 μm/s or ∼ 0.751 mm/day) and their anterograde bias match well with the results of radiolabeling experiments^2, 3^, suggesting that they represent the same phenomenon. We could also detect a second rate-class of spectrin, which moved at ∼0.84 μm/s, paused frequently (consistent with a “Stop and Go” model) and had a fainter signal compared to the slower fraction (Fig. 3E, Fig. S7). This might correspond to a fraction of spectrin that was shown to move at 50 mm/day by radiolabeling, similar to mitochondrial proteins^6^. Thus, spectrin travels in at least two distinct rates, with the slower one being nearly as processive (i.e., runs without stopping) as the faster one.

**Figure 3:**
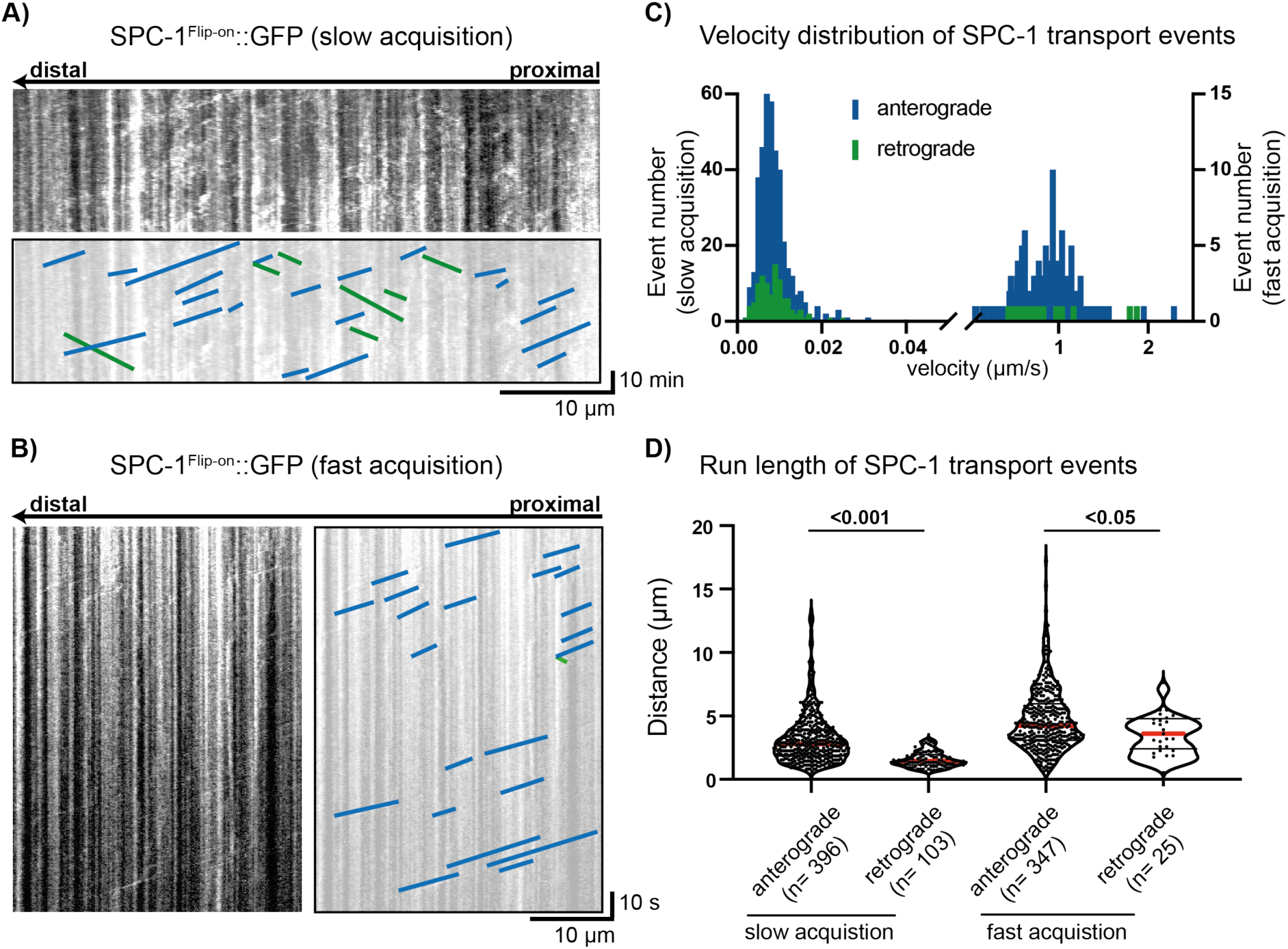
Slow transport of endogenous spectrin is processive. A. Representative kymograph showing slow and processive movement of spectrin (SPC-1^Flip-on^::GFP, 8-14 hrs after induction) in the axon. Acquisition rate: 30 s/ frame. Bottom panel highlights movement events in blue (anterograde) and green (retrograde). B. Representative kymograph showing intermittent fast movement of spectrin in the axon. Acquisition rate: 0.3 s / frame. C. Velocity distribution of SPC-1^Flip-On^::GFP showing two distinct transport rates. D. Run length analysis of SPC-1^Flip-on^::GFP showing that the slow movement of spectrin is processive. Data in B+D were collected from n= 13 (slow acquisition) and n= 20 (fast acquisition) animals. Statistical analysis is based on a Kruskal-Wallis test, followed by Dunńs post test for multiple comparisons.

We next examined the SPC-1^Flip-On^::GFP probe in combination with UNC-76 single cell degradation (*unc-76 scKD*) or the kinesin-1 allele *unc-116(e2310),* which contains a transposon insertion at its C-terminus cargo binding region^25^ that is predicted to disrupt the interaction between UNC-76/FEZ1 and kinesin^20^.

We found that at early timepoints after heat shock, the distribution of newly synthesized SPC-1^Flip-on^::GFP was shifted towards the cell body in both *unc-116(e2310)* and *unc-76 scKD* mutants, consistent with transport defects (Fig. S8A). Direct visualization of slow transport over a one-hour imaging window showed almost no effect on retrograde transport but a drastically reduced number of movement events in the anterograde direction (Fig. S8B, C). The number of faster transport events in both mutants was also reduced (Fig. 8D, E). These results indicate that UNC-76 is required for the axonal transport of spectrin.

To test whether UNC-76/FEZ1 and UNC-69/SCOC recruit spectrin to kinesin for transport, we first tested whether FEZ1, SCOC and SPTBN1 (human βII spectrin) co-localize when transfected into COS7 cells. Although endogenous FEZ1 localizes to puncta^22^, overexpressed SNAP::FEZ1 and HALO::SCOC were diffuse, while SPTNB1::GFP was enriched near the plasma membrane (Fig S9). However, we also observed clear events, albeit infrequent, where the three proteins colocalized and co-migrated (Fig S9B, C). Consistent with these results, we co-immunoprecipitated HALO::SCOC and mCherry::FEZ1 with SPTBN1::GFP from transfected HEK293T lysates (Fig S10). These results suggest that FEZ1 and SCOC can form a transport packet that contains spectrin.

To test *in vivo* whether UNC-76/FEZ1 functions as a cargo adaptor for spectrin, we asked whether it can be replaced by artificially linking spectrin to kinesin. We generated a construct containing kinesin light chain 2 fused to anti-GFP nanobody and expressed it in *unc-76 scKD; SPC-1::GFP* animals (Fig. 4A). This construct significantly improved spectrin distribution, suggesting that the function of UNC-76/FEZ1 is to link spectrin to kinesin (Fig. 4B). Finally, we asked whether UNC-76 could instruct excessive spectrin transport when overexpressed. Overexpressed RFP::UNC-76 accumulated at the tip of the DA9 axon (Fig. 4C_(i)_). RFP::UNC-76 generated an ectopic accumulation of spectrin at the tip of the axon, with spectrin levels there correlating to UNC-76 levels (Fig 4C_(i,ii)_). In contrast, a control kinesin-1 cargo, MIRO-1::GFP, did not accumulate at the distal axon (Fig 4C_(i,iii)_). Mutations in *unc-69* and *unc-116* eliminated the distal RFP::UNC-76 and SPC-1::GFP accumulations (Fig. S11). Furthermore, in *unc-116* mutants, RFP::UNC-76 and SPC-1::GFP co-localized in punctate structures in the cell body (Figure 4D_(iv, v)_), consistent with an assembled transport particle which fails to move. Together, these results strongly support the conclusion that UNC-76/FEZ1 and UNC-69/SCOC are a cargo adaptor for spectrin transport and can instruct the distribution of axonal spectrin.

**Figure 4:**
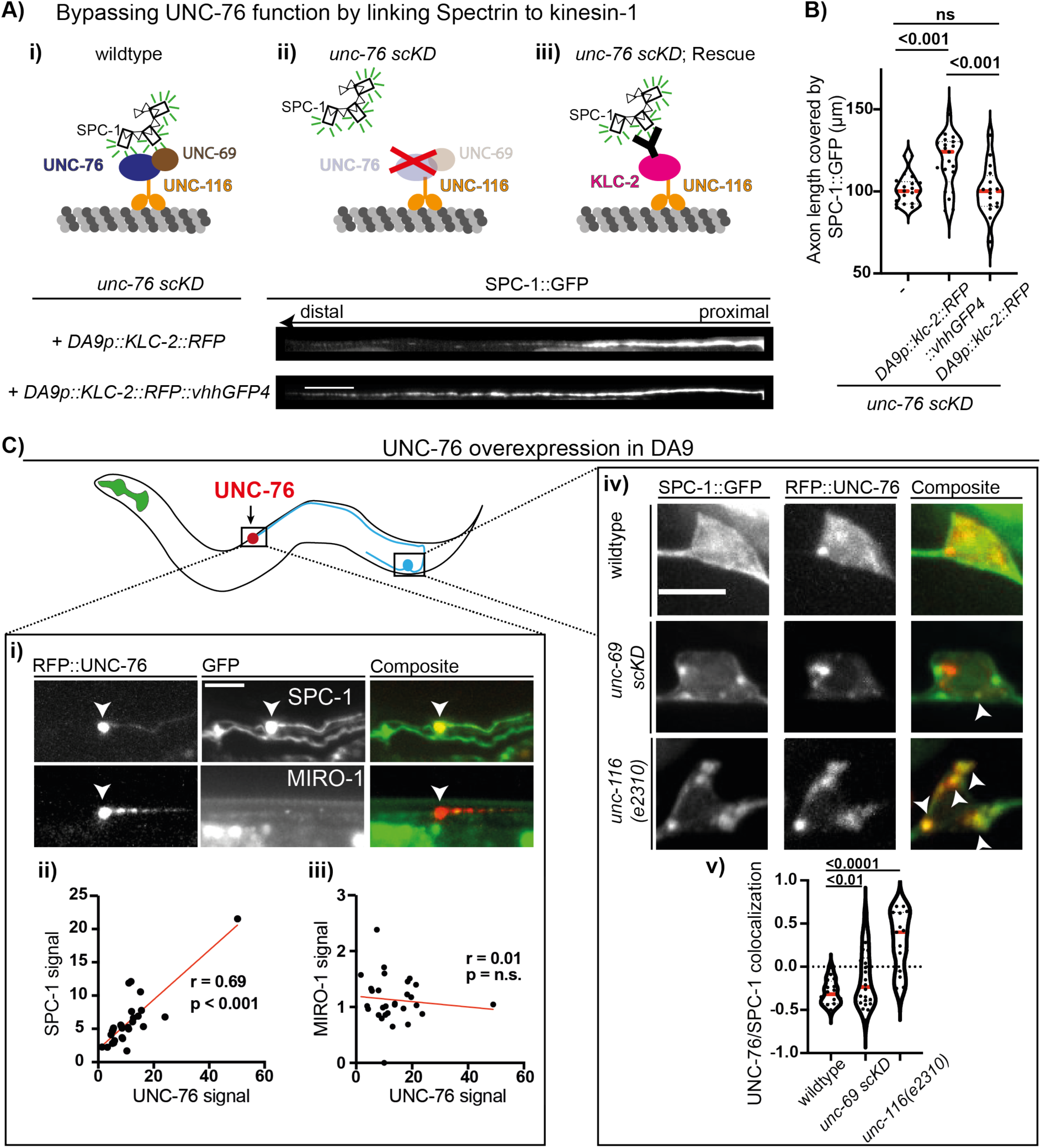
UNC-76/FEZ1 links spectrin to kinesin-1. A. Rescue of SPC-1 localization to the distal axon in *unc-76 scKD* by directly linking α-spectrin to kinesin-1. Top panels show the experimental design. SPC-1::GFP was expressed in an *unc-76 scKD* animals with a control KLC-2::RFP (ii) or KLC-2::RFP::vhhGFP4 (iii). Lower panels show representative images of SPC-1::GFP in control (ii) and rescue conditions (iii). Scale: 10 μm. B. Quantification of results from A. n = 16-21 per genotype, 1-way ANOVA followed by Tukeýs post-test for multiple comparisons. C. Overexpression of UNC-76 instructs spectrin localization to the tip of the axon. i) Representative images of overexpressed RFP::UNC-76 with SPC-1::GFP or with a control kinesin-1 cargo (MIRO-1::GFP labels mitochondria) at the axon tip. Scale:10 μm. Note accumulation of spectrin, but not mitochondria in the tip of the axon. Bottom: linear regression of RFP::UNC-76 signal intensity and SPC-1::GFP (ii, n = 28) or MIRO-1::GFP (iii, n = 29) signal intensity. iv) SPC-1 colocalizes with RFP::UNC-76 (indicated by white arrowheads) in the cell bodies of *unc-116* mutants and *unc-69 scKD*. v) Quantification of the colocalization in (iv) using Pearson correlation coefficient.

Here we visualized the slow axonal transport of spectrin from the neuronal cell body to hot spots in the spectrin lattice and determined that hotspots support MPS expansion at the rate of axon stretch-growth. Our results reveal that slow transport of spectrin – and potentially other cargo – is processive, raising important mechanistic questions. To begin answering these questions, we identified a complex of UNC-116/UNC-76/UNC-69 that mediates spectrin transport.

## Acknowledgements

The authors would like to thank the Caenorhabditis Genetic Center (CGC) for sharing strains. This project is funded by the NIH grants R35-GM131744 (SY) and AG062210 (SMF). OG is supported by a Walter-Benjamin Scholarship funded by the Deutsche Forschungsgemeinschaft (DFG, German Research Foundation) -Project# 465611822. We would like to thank the entire Yogev and Hammarlund labs for technical assistance, feedback and discussions.

## Author contribution

OG, GS and SY designed, performed and analyzed all nematode related experiments, designed the figures and wrote the manuscript. SS and JH analyzed the hotspot patterning. PML, CL and SMF performed the IP from HEK293T cells. JP transfected and imaged SPTBN1/FEZ1/SCOC in Cos7 cells. YW and MH generated the 7xspGFP11::MIRO-1 knock in strain.

## Supplementary Figures

**Fig. S1:**
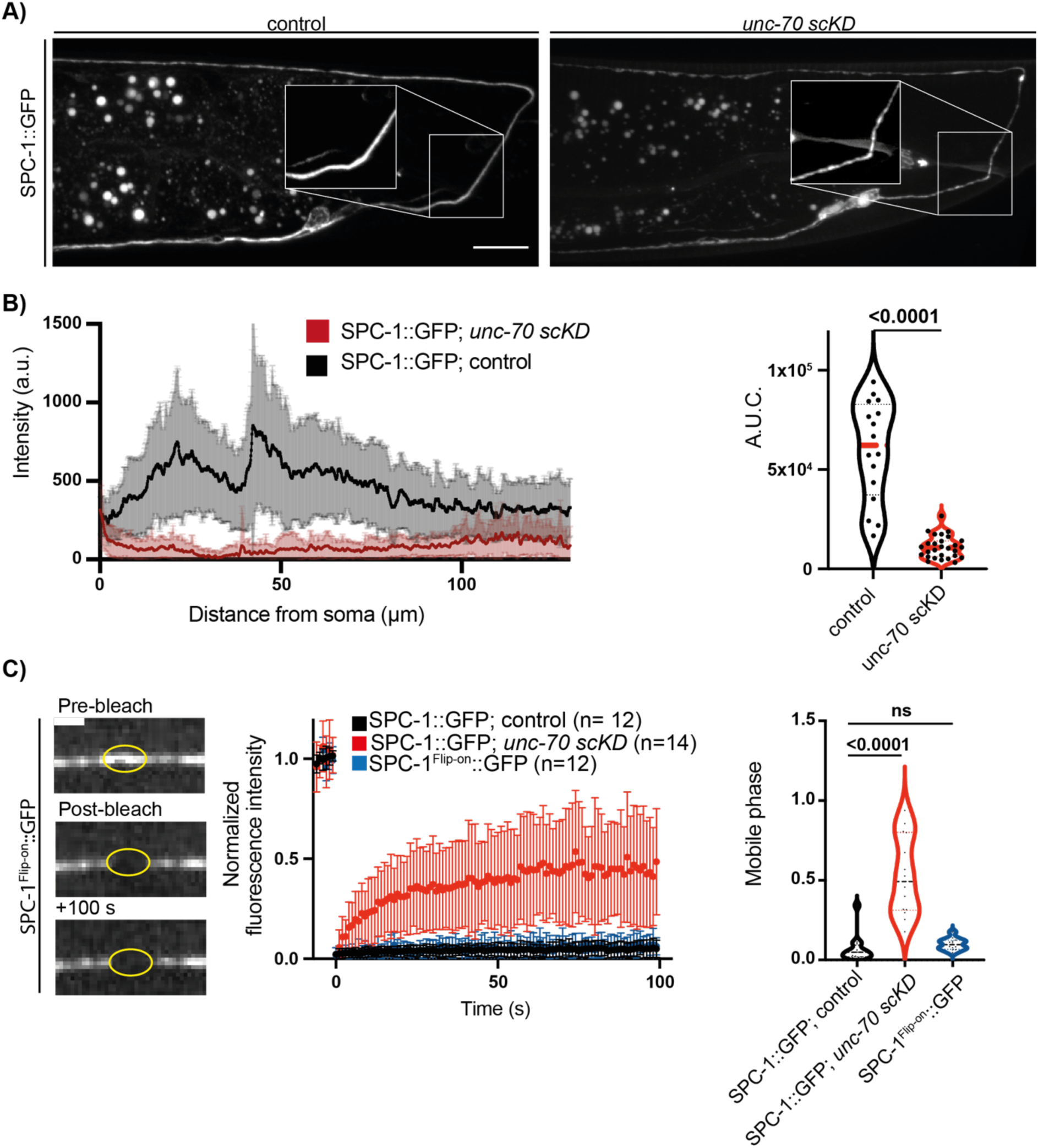
SPC-1::GFP integrates well into the MPS lattice. A) A cell specific knock down of neuronal ß-spectrin (UNC-70) reduces the axonal localization of SPC-1::GFP. Single cell knock down (scKD) was achieved by tagging the genes with an AID degron, expressing TIR1 in DA9 and growing animals on Auxin-containing NGM plates. Control animals do not express TIR1. Boxed region highlights the change in signal distribution in the proximal axon / commissure upon knock down. Scale: 10 μm. B) SPC-1::GFP signal intensity in the first 130 μm of the axon were measured to determine the area under the curve and compared between control and *unc-70 scKD*. Statistical comparison is based on an unpaired t-test. C) The mobility of SPC-1::GFP and SPC-1^Flip-on^::GFP (8 hrs after induction) was tested using FRAP. Hardly any recovery is observed, suggesting integration into the MPS lattice. *unc-70 scKD* increases SPC-1::GFP recovery, confirming FRAP as an assay for the integration of SPC-1::GFP into the MPS. Statistical comparison is based on a Kruskal-Wallis test followed by a Dunńs test for multiple comparisons.

**Fig. S2:**
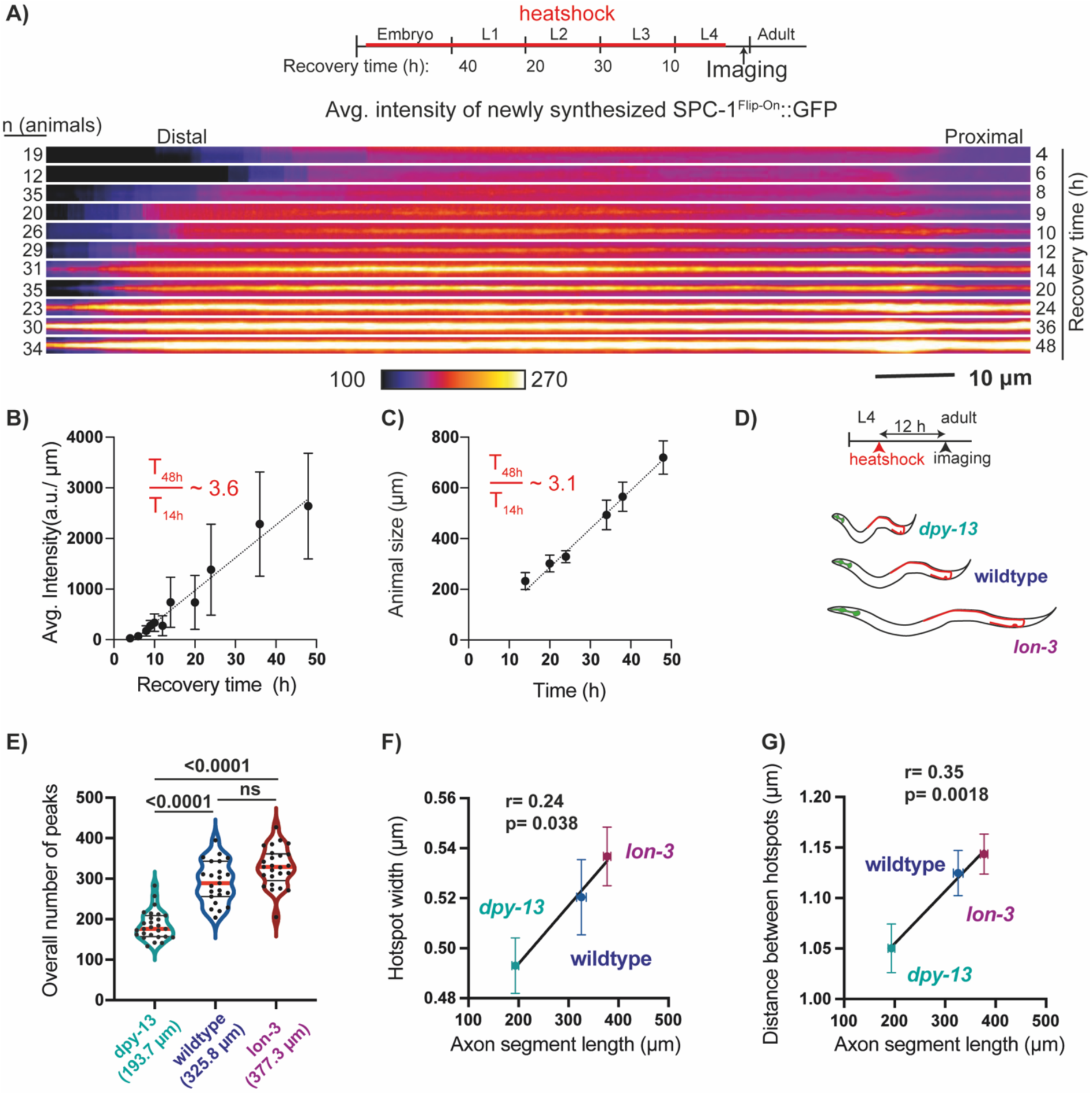
Correlation of MPS expansion from hotspots with axon stretch-growth. A. timeline of experimental scheme (top): animals were heat shocked at indicated time points and left to recover for the indicated amount of time until imaging as late L4 larvae. Bottom panel shows pseudo-colored fluorescence intensity profiles of SPC-1^Flip-on^::GFP, averaged from 12-35 animals per timepoint. B. Quantification of data from A. Signal intensity increases ∼3.6 fold between 14 and 48 hrs. Note that the signal increase is linear, suggesting that a steady-state of spectrin incorporation and removal from the MPS has not been reached (in case of a steady state we would expect the intensity profile to plateau). C. Animal growth along the A-P axis during the time of the experiment. Notice that the increase in animal growth (∼3.1 fold) and therefore DA9 stretch-growth is similar to the increase in SPC-1^Flip-on^::GFP intensity. Error bars in B and C are for s.d. E, F, G. total number (E), width (F) and distance between peaks (G) of SPC-1^Flip-on^::GFP hotspots at 12 hours after induction in *lon-3* (n= 25), *dpy-13* (n= 27) and wildtype (n= 23) animals. E. Quantification of the number of hotspots per genotype in the entire segment analyzed. Note that this segment is significantly shorter in *dpy-13* mutants. Statistical comparison is based on a Kruskal-Wallis test followed by Dunńs post-test for multiple comparisons. Correlation of hotspot width (F) and the distance between hotspots (G) with axon segment length by Pearson. R and p-value are indicated. Error bars in F, G are SEM.

**Fig. S3:**
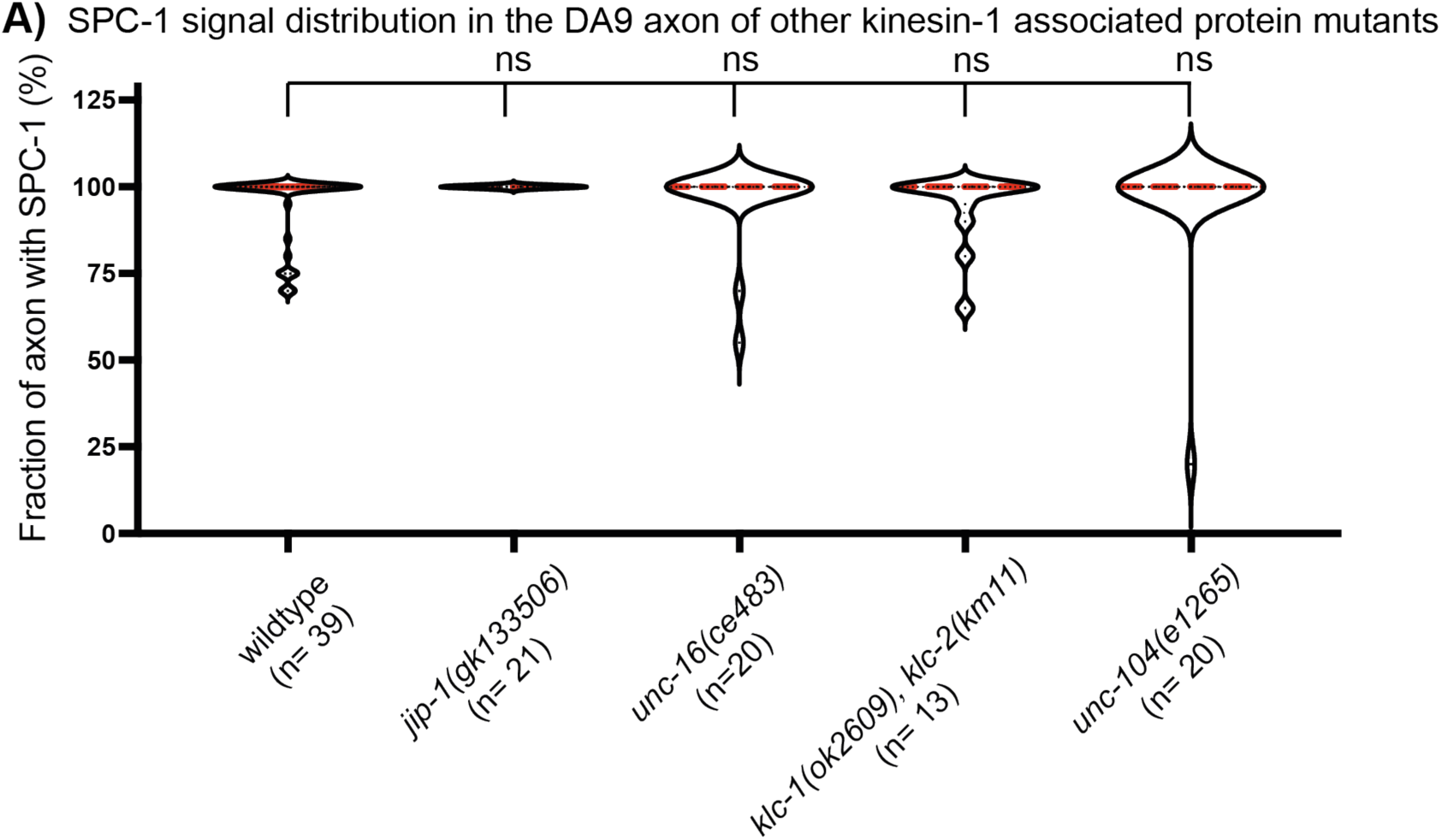
Mutations in various kinesin-1 adaptors do not affect spectrin distribution. A. Quantification of the axonal fraction that contained SPC-1::GFP signal (see material and methods for details) in wildtype and indicated genotypes. Red line and faint black lines in violin indicate mean and 25^th^/75^th^ percentiles, respectively. n indicated on graph.

**Fig. S4:**
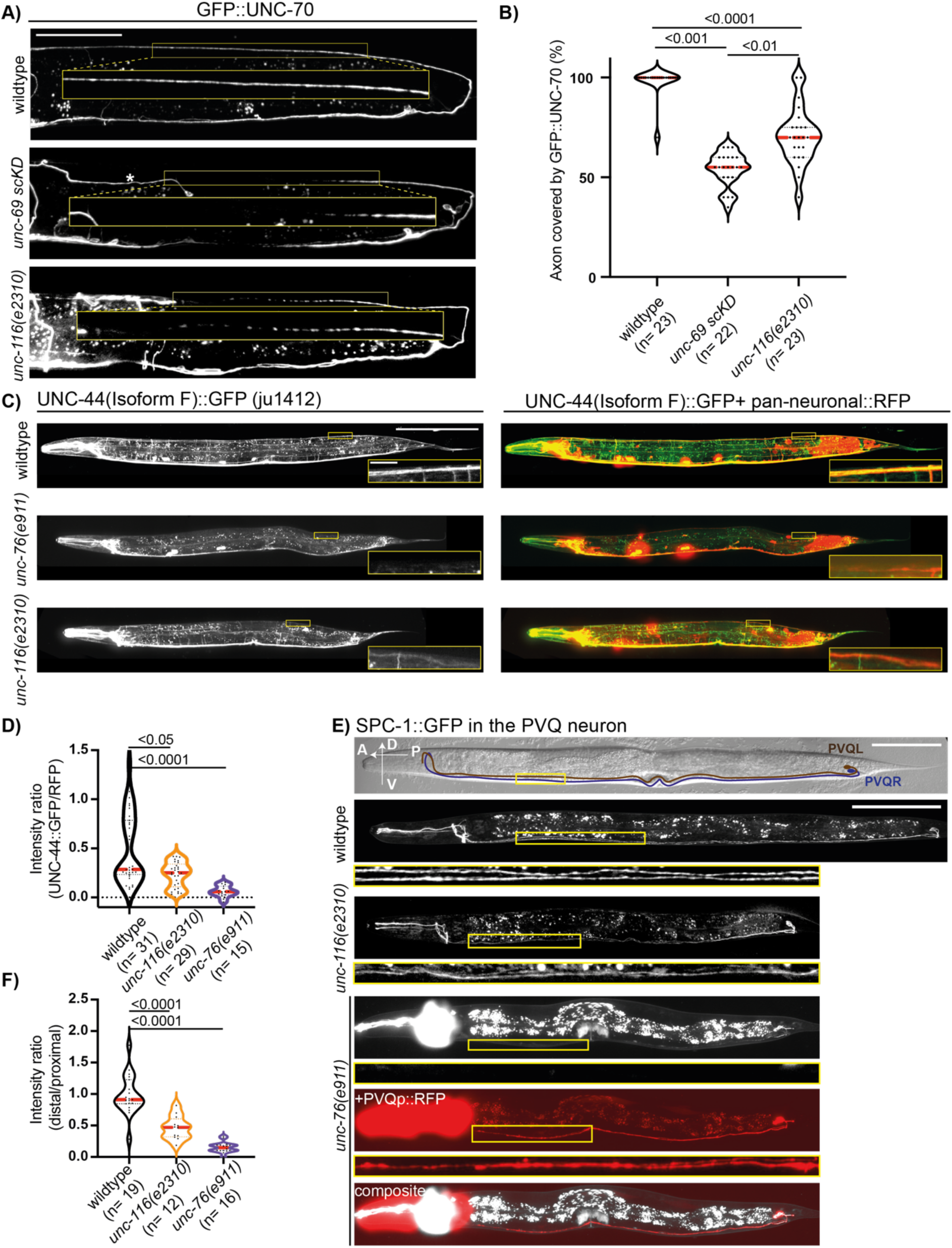
*unc-76, unc-69* and *unc-116* are required for the distribution of MPS proteins in different neurons. A. UNC-70/β spectrin was endogenously tagged with 7xspGFP11 and visualized in DA9 with a *mig-13* promoter driving spGFP1-10. GFP::UNC-70 is lost from the distal axon in *unc-116 (e2310)* and *unc-69 scKD* backgrounds. Scale: 50 μm. B. Quantification of A, Kruskal-Wallis test followed by Dunńs test for multiple comparisons. C. Isoform F of UNC-44/Ankyrin was visualized in the entire animal by a GFP inserted in this specific isoform (*ju1412),* along with pan-neuronal RFP (*Prab-3::RFP). unc-76(e911)* and *unc-116(e2310)* mutants show loss of UNC-44, including in regions where the DA9 axon is present (shown by RFP in insets). Scale: 100 μm. D. Quantification of C. UNC-44(isoform F)::GFP signal was normalized to the RFP signal in the dorsal nerve cord to determine the loss of UNC-44(isoform F) from neurites of the dorsal nerve cord. Statistical comparison is based on a Kruskal-Wallis test followed by an uncorrected Dunńs test. E. SPC-1::GFP localization is disrupted in PVQ neurons of *unc-76(e911)* and *unc-116(e2310)* animals. Schematic of PVQR and PVQL neurons is shown in the top panel. Since the signal intensity in the distal PVQ neurons of *unc-76 (e911)* mutants is often undetectable, those animals also expressed a cytosolic RFP (*Psra-6::tagRFP*) to show that the neurons were still present. Yellow boxed area below the main figures show a magnification in the distal axon. Scale:100 μm. SPC-1::GFP distribution is quantified in F as a ratio of signal intensity in the distal 50% of the axon to the proximal 50%. Statistical comparison is based on an ordinary 1-way ANOVA, followed by Tukeýs post-test for multiple comparison.

**Fig. S5:**
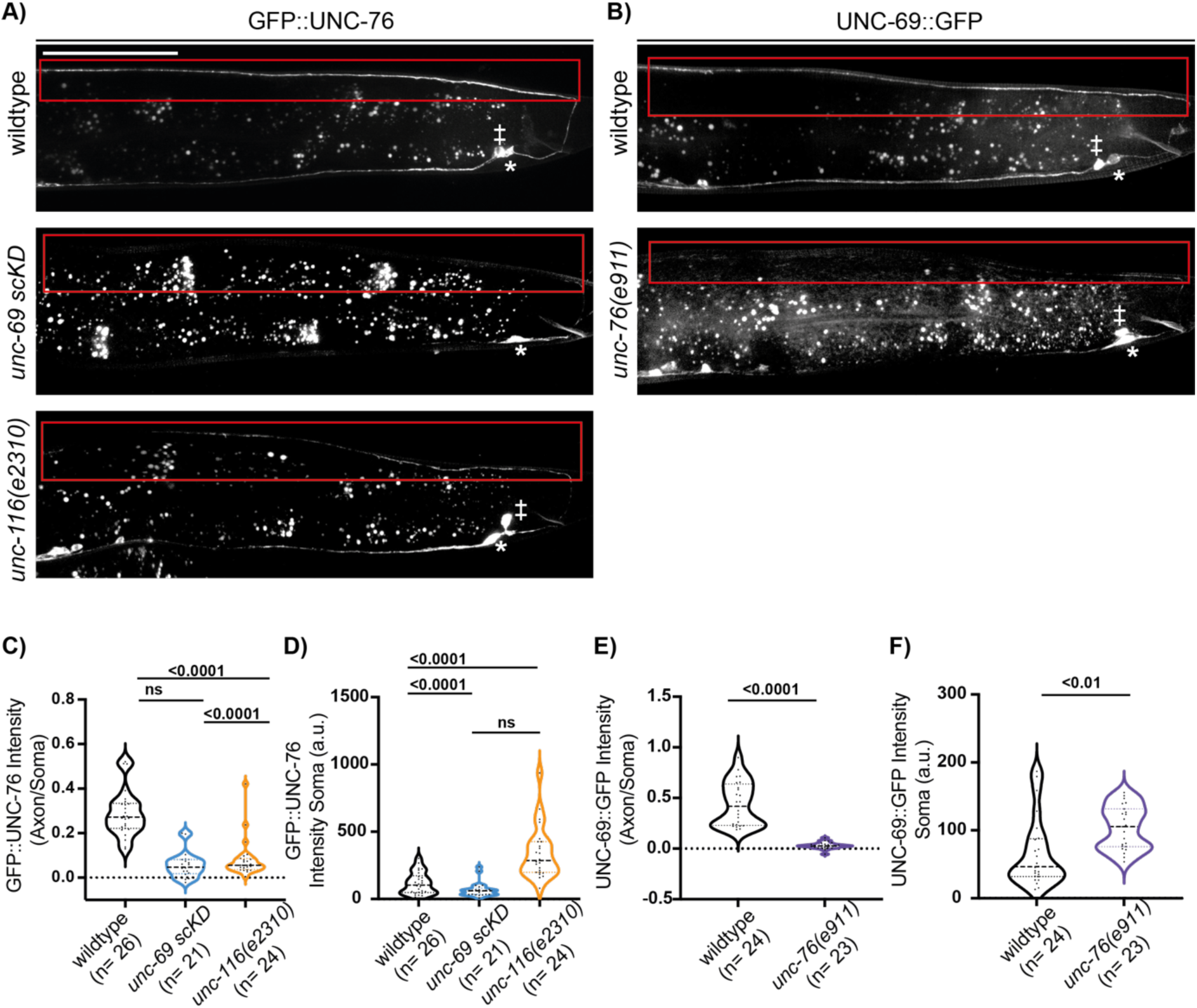
UNC-76 and UNC-69 depend on each other and on UNC-116 to localize robustly throughout the axon. A. UNC-76, tagged endogenously with 10xspGFP11 and visualized with a DA9 specific spGFP1-10, localizes throughout the axon. Axonal levels of UNC-76 are drastically reduced in *unc-116 (e2310)* mutants and *unc-69 scKD.* B. UNC-69, tagged endogenously with 7xspGFP11 and visualized with a DA9 specific spGFP1-10, localizes throughout the axon. UNC-69 levels are drastically reduced in *unc-76 (e911)* mutants. Boxed regions in A and B indicate the dorsal part of the DA9 axon. Asterisks mark the DA9 cell body. The cell body of VA12, which is also labelled by the *mig-13* promoter, is indicated with a ‡. Scale: 50 μm. C+D. Quantification of the UNC-GFP::76 signal distribution as a ratio of the average signal intensity in the axon to the average signal intensity in the soma (C) or as the average signal intensity in the soma (D) of the strains shown in A. Statistical comparison in both data sets is based on a Kruskal-Wallis test, followed by Dunńs post test for multiple comparisons. E+F. Quantification of the UNC-69::GFP signal distribution as in C+D but with the strains shown in B. Statistical comparison in E+F is based on a Mann-Whitney test.

**Fig. S6:**
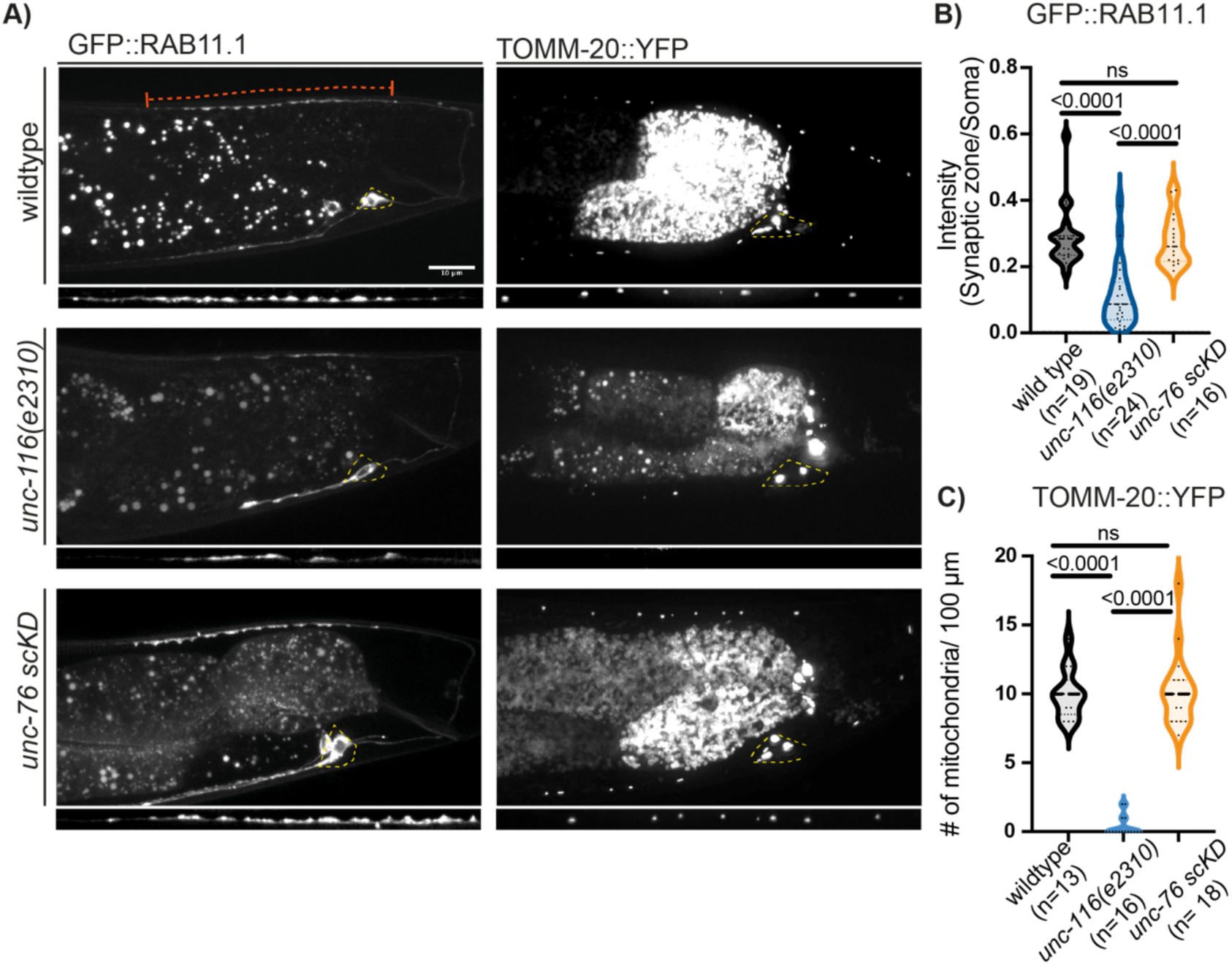
*unc-76* is not required for the distribution of kinesin-1 cargos mitochondria and RAB11.1 vesicles. A. The localization of mitochondria and RAB11.1 positive vesicles depends on UNC-116 but not UNC-76. Images show nematodes expressing GFP::RAB-11.1 (left panels) or TOMM-20::YFP to label mitochondria (right panels) in the DA9 neuron of wildtype, *unc-116(e2310)* or *unc-76 scKD* backgrounds. The axonal signal (indicated by an orange line in top left image) is magnified below each image. Encircled areas indicate the cell body. Note that both markers in a wildtype or *unc-76scKD* background are robustly localized to axon. Scale: 10 μm. B. Quantification of the signal enrichment of GFP::RAB-11.1 in the synaptic zone from strains shown in A, based on the ratio of the signal intensity at the synaptic zone to the signal intensity in the soma. Statistical comparison is based on a Kruskal-Wallis test followed by Dunńs post-test for multiple comparisons. C. Quantification of the mitochondria distribution in the first 100 μm of the DA9 axon on the dorsal side of the nematode Statistical comparison is based on a Kruskal-Wallis test followed by Dunńs post-test for multiple comparisons.

**Fig. S7:**
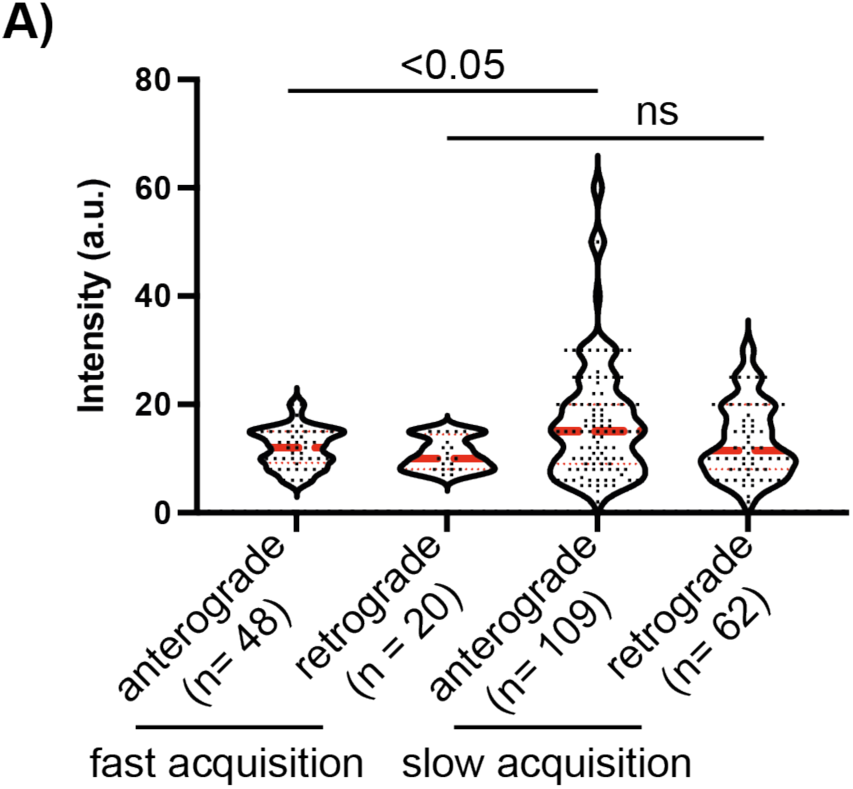
Comparison of SPC-1^Flip-on^::GFP intensity in the two transport populations. A. Signal intensities of single movement events (after background subtraction) were quantified from the kymographs derived from the time lapse experiments of SPC-1^Flip-on^::GFP (see Fig.3) at slow or fast (intermediate) acquisition speed. Statistical comparison is based on a Kruskal-Wallis-test followed by an uncorrected Dunńs post test for multiple comparisons.

**Fig S8:**
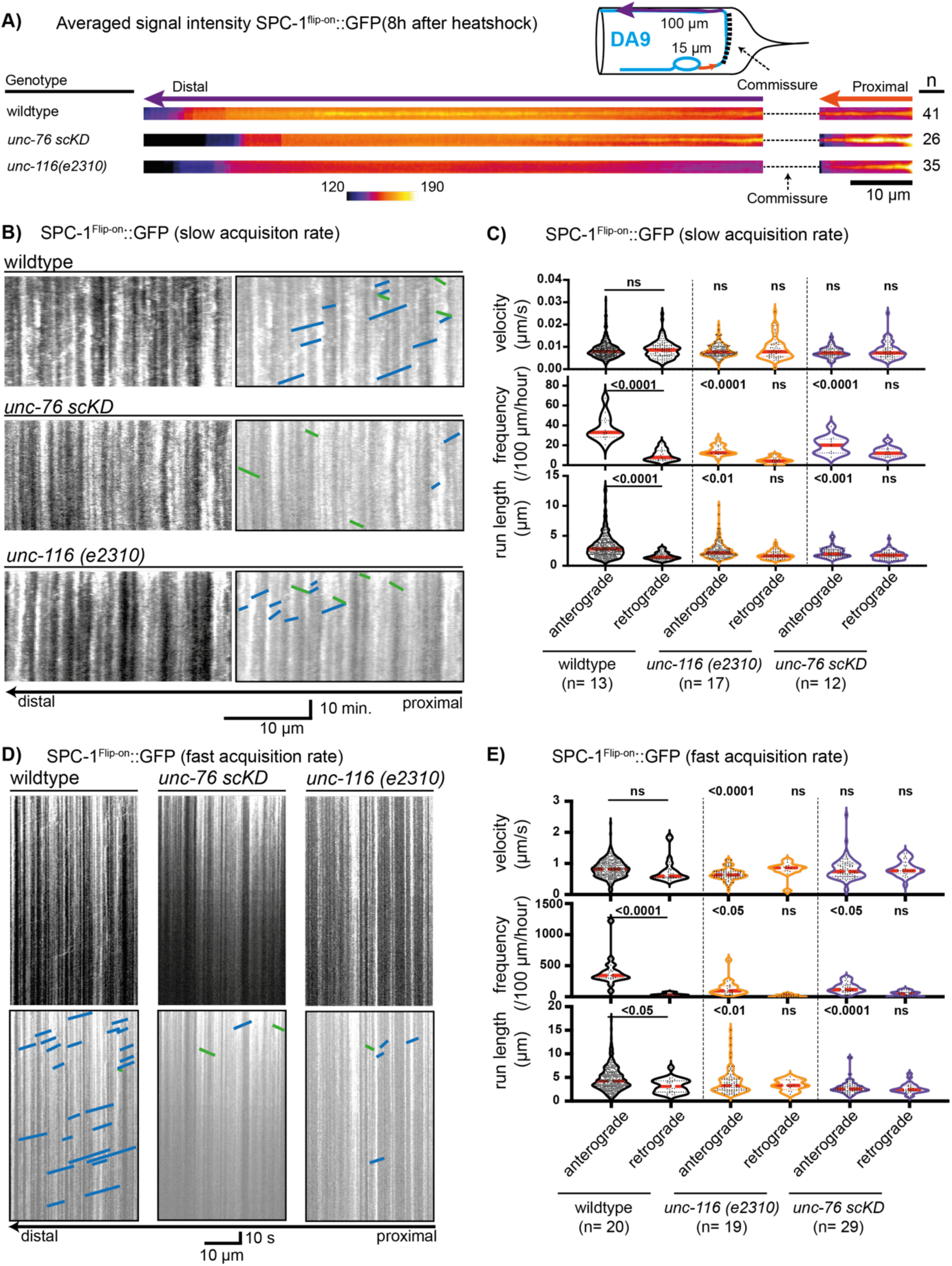
spectrin transport depends on UNC-116, UNC-76 and UNC-69. A. Averaged pseudo-colored line scans of SPC-1^Flip-on^::GFP at 8 hrs after induction from animals of the indicated genotypes. Note proximal shift in the distribution of newly synthesized spectrin in *unc-116* mutants and *unc-76 scKD.* B, C. Representative kymographs (B) and quantifications (C) of SPC-1^Flip-on^::GFP (8-14 hrs after induction) transport parameters under slow acquisitions in the indicated genotypes. Note drastic reductio of movement events in *unc-116 (e2310)* mutants and *unc-76 scKD* animals. n = 12-17 animals per genotype, ordinary 1-way ANOVA followed by Tukeýs post-test for multiple comparisons (run length) or a non-parametric 1-way ANOVA followed by Dunńs post-test for multiple comparisons (speed and frequency), p values indicated on plot. D, E. Similar to B and C, except for the use of faster acquisition rates, which reveals the SPC-1^Flip-on^::GFP moving at ∼0.8 μm/s. n = 19- 29 animals per genotype, non-parametric 1-way ANOVA followed by Dunńs post-test for multiple comparisons, p values indicated on graph.

**Fig. S9:**
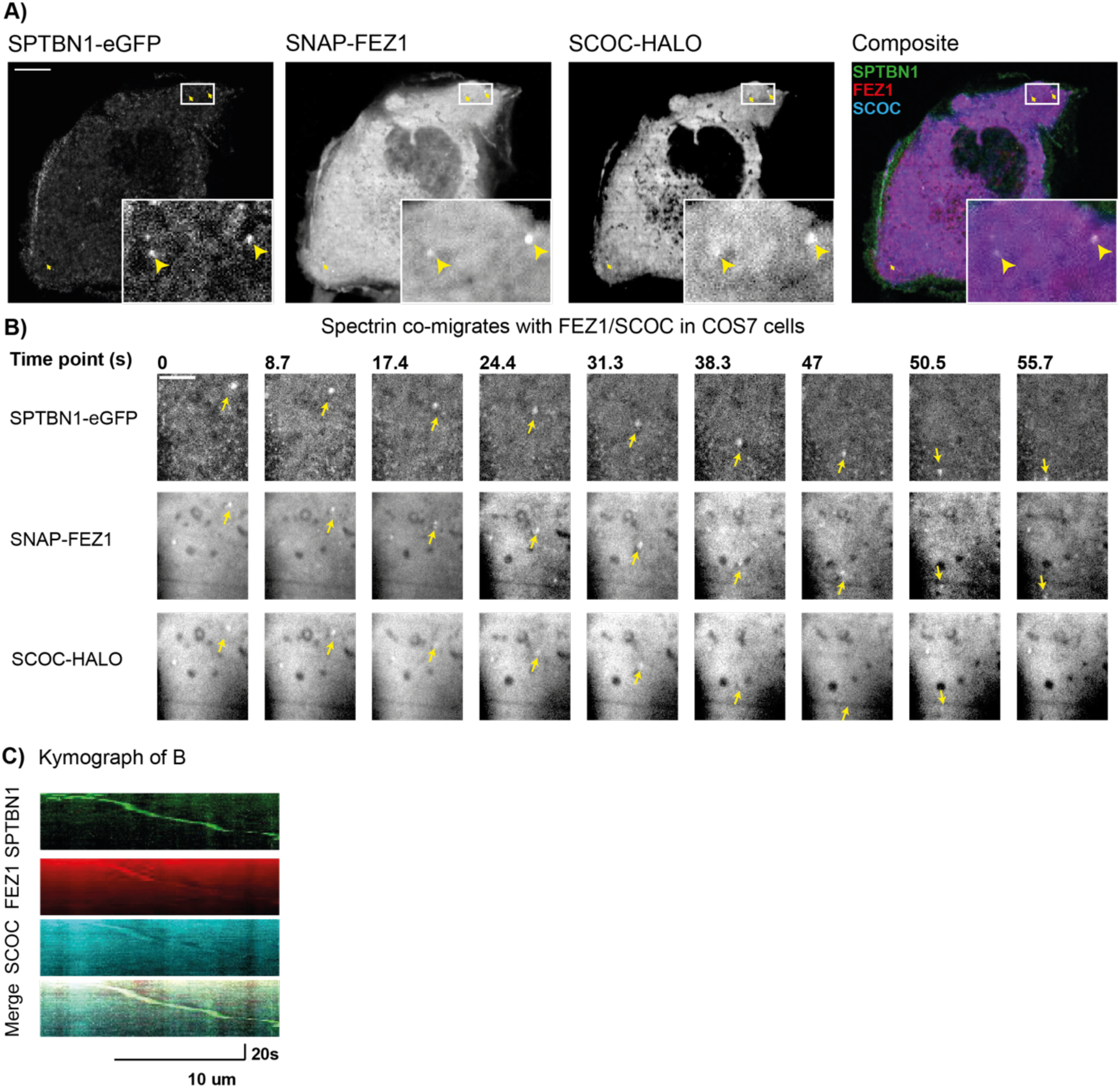
Spectrin is in a transport packet with FEZ1/SCOC in COS7 cells. A. SPTBN1 is mostly in the membrane but can also be observed in distinct puncta which contain FEZ1 and SCOC. COS7 cells were transfected with SPTBN1-eGFP, SNAP-FEZ1 and SCOC-HALO, incubated with SNAP and HALO tag ligand conjugated fluorophores and imaged on a spinning-disc confocal microscope at 63x. Arrows indicate co-localization events of all 3 proteins. Scale bar equals 10 μm. B. SPTBN1 co-migrates with FEZ1 and SCOC in distinct puncta. Multiple images were acquired as in A of both a single focal plane and as a time-lapse experiment with an interval of 1.74 s between each timepoint. An example of a co-movement event of all three proteins is shown. Timepoint 0 marks the first observation event of this SPTBN1/SCOC/FEZ containing puncta. Scale: 5 μm. C. Kymograph of the moving puncta from B for all three fluorescent channels and as a merged image.

**Fig. S10:**
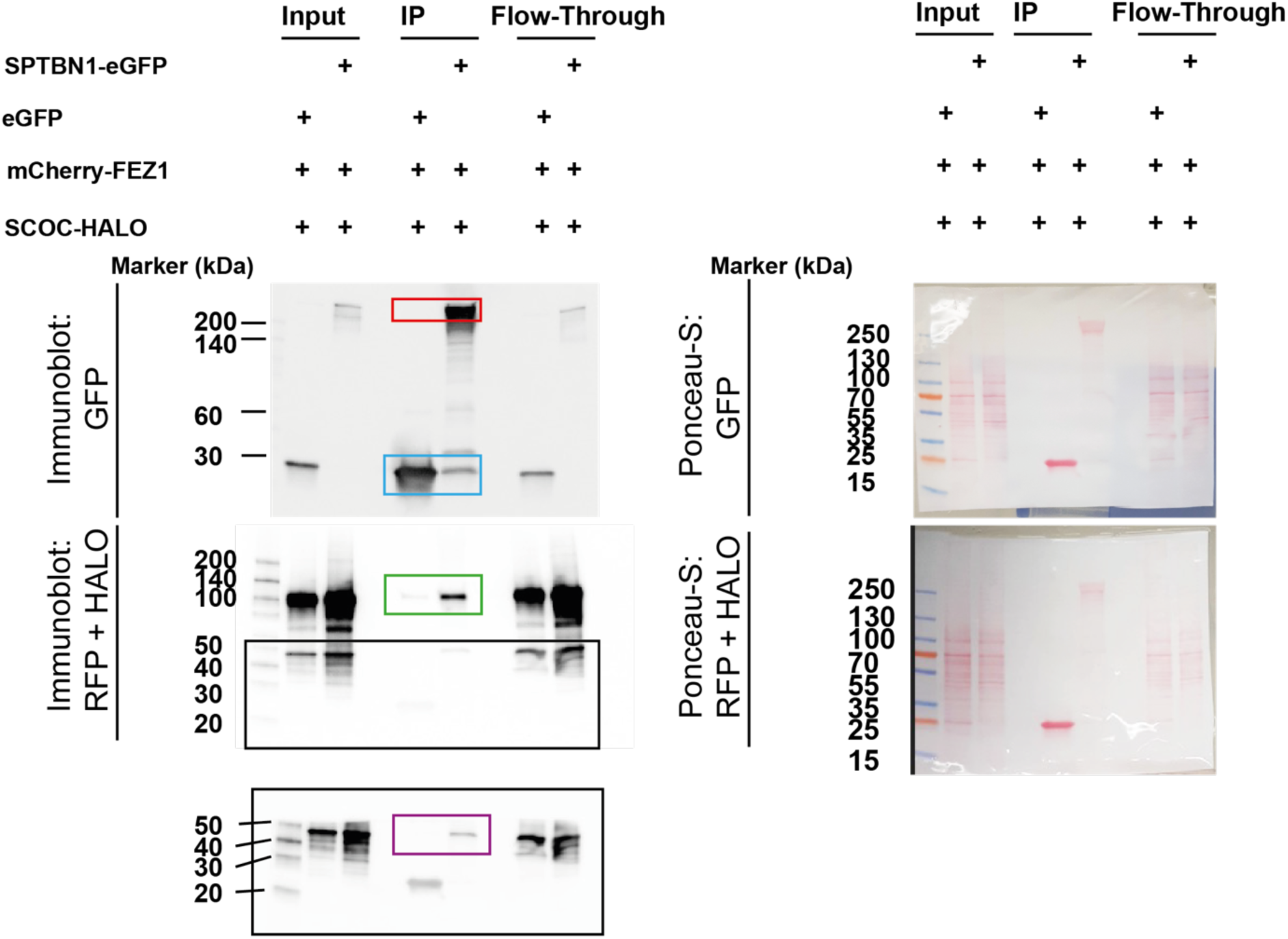
Spectrin co-immunoprecipitates with FEZ1/SCOC from Hek293T cell lysates. HEK293T cells were transfected with SCOC-HALO, mCherry-FEZ1 and either SPTBN-1-eGFP or eGFP (control). Lysates were incubated with GFP nanobody coupled beads for pull down. Left panels: The upper membrane was incubated with a GFP antibody, the middle membrane co-incubated with an RFP and HALO antibodies (see Material and Methods for details). Following second antibody incubation and detection with ECL solution, the middle membrane was cut (indicated by the black box) and the lowest membrane was incubated with Femto-ECL solution to better detect the HALO-SCOC band. Boxed colored regions indicate the expected sizes for SPTBN-1 (red), eGFP (blue), FEZ1 (green) and SCOC (purple).

**Fig. S11:**
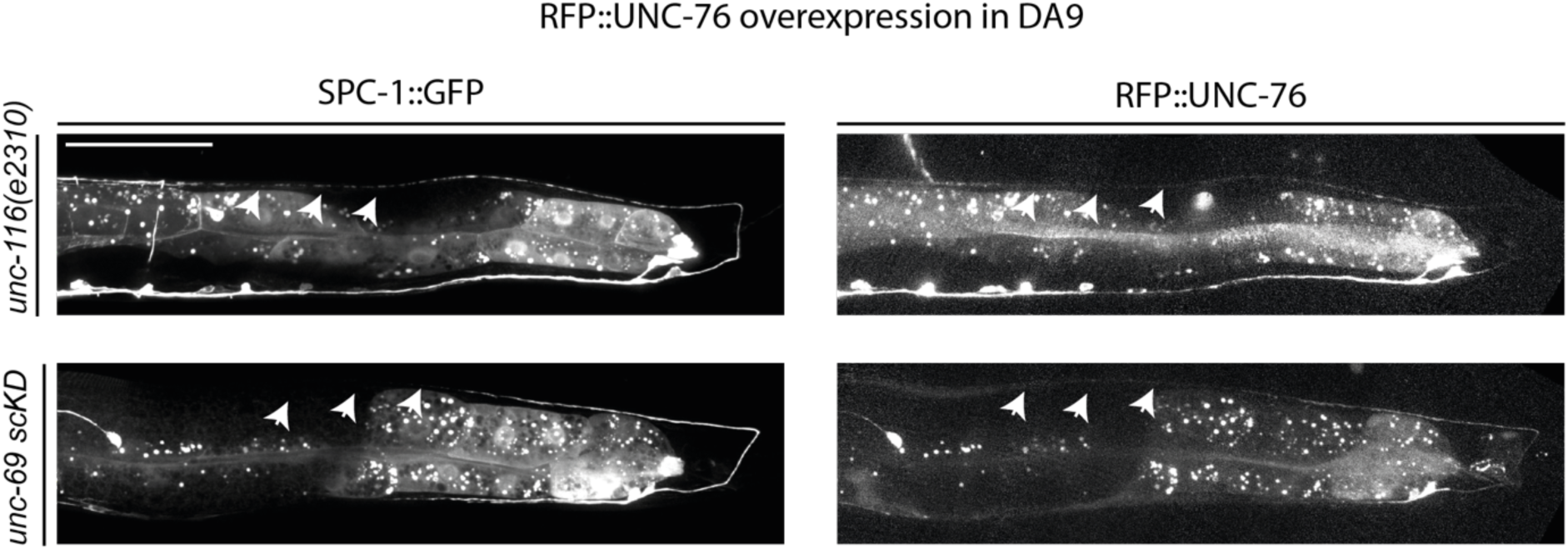
The distal co-accumulation of SPC-1 and overexpressed RFP::UNC-76 depends on UNC-116 and UNC-69. RFP::UNC-76 was overexpressed in *unc-116 (e2310)* mutants or *unc-69 scKD* animals harboring SPC-1::GFP. In the mutants (compare to wildtype in Fig. 4), SPC-1::GFP and RFP::UNC-76 signals become very dim or absent towards the distal end of the DA9 axon (indicated by white arrows). Scale: 50 μm.

## Material and Methods

### Strains and maintenance

Nematode strains were cultured on nematode growth medium plates that were seeded with OP50 bacteria. Animals were grown at 20°C unless indicated otherwise.

### Strain collection

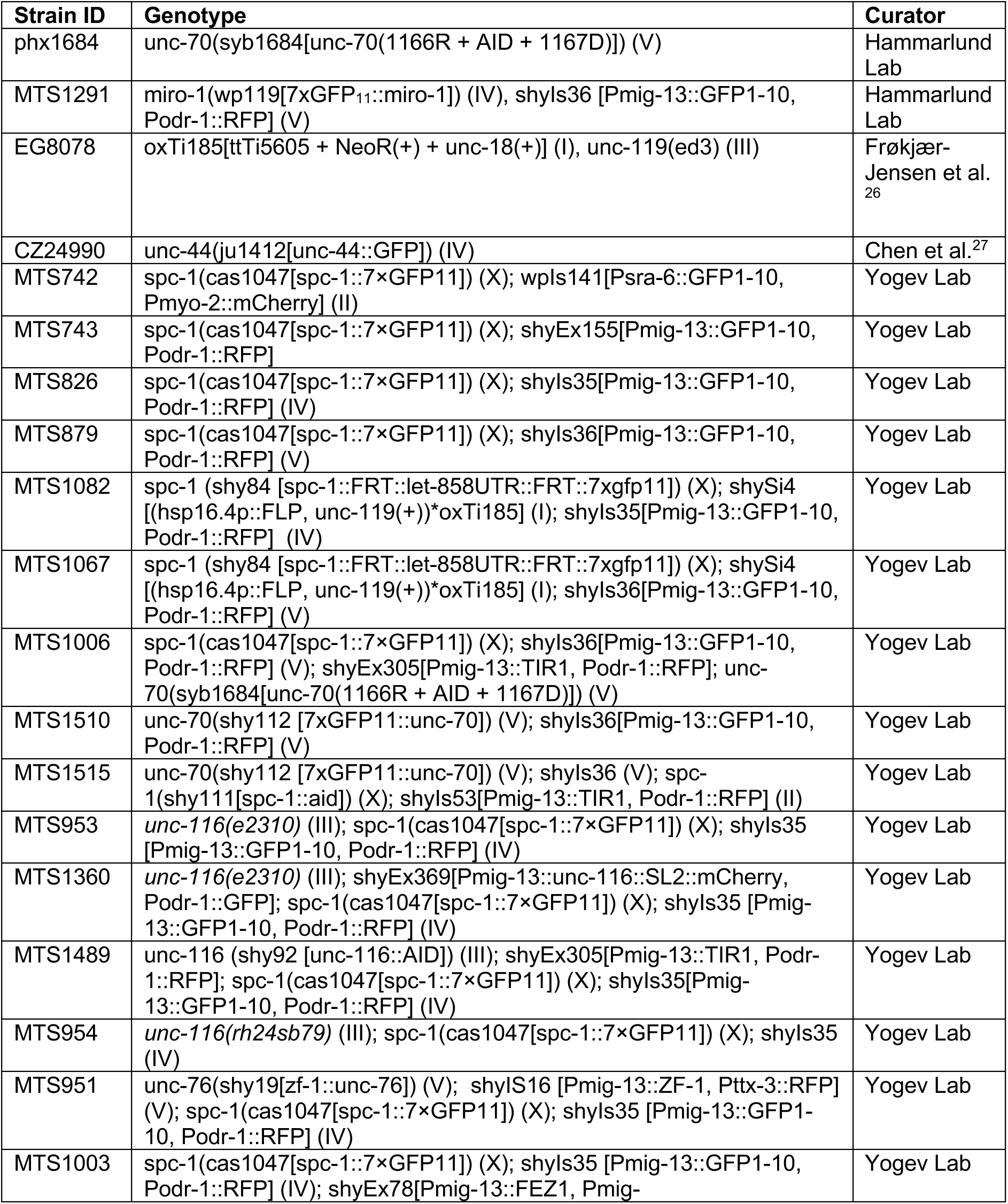
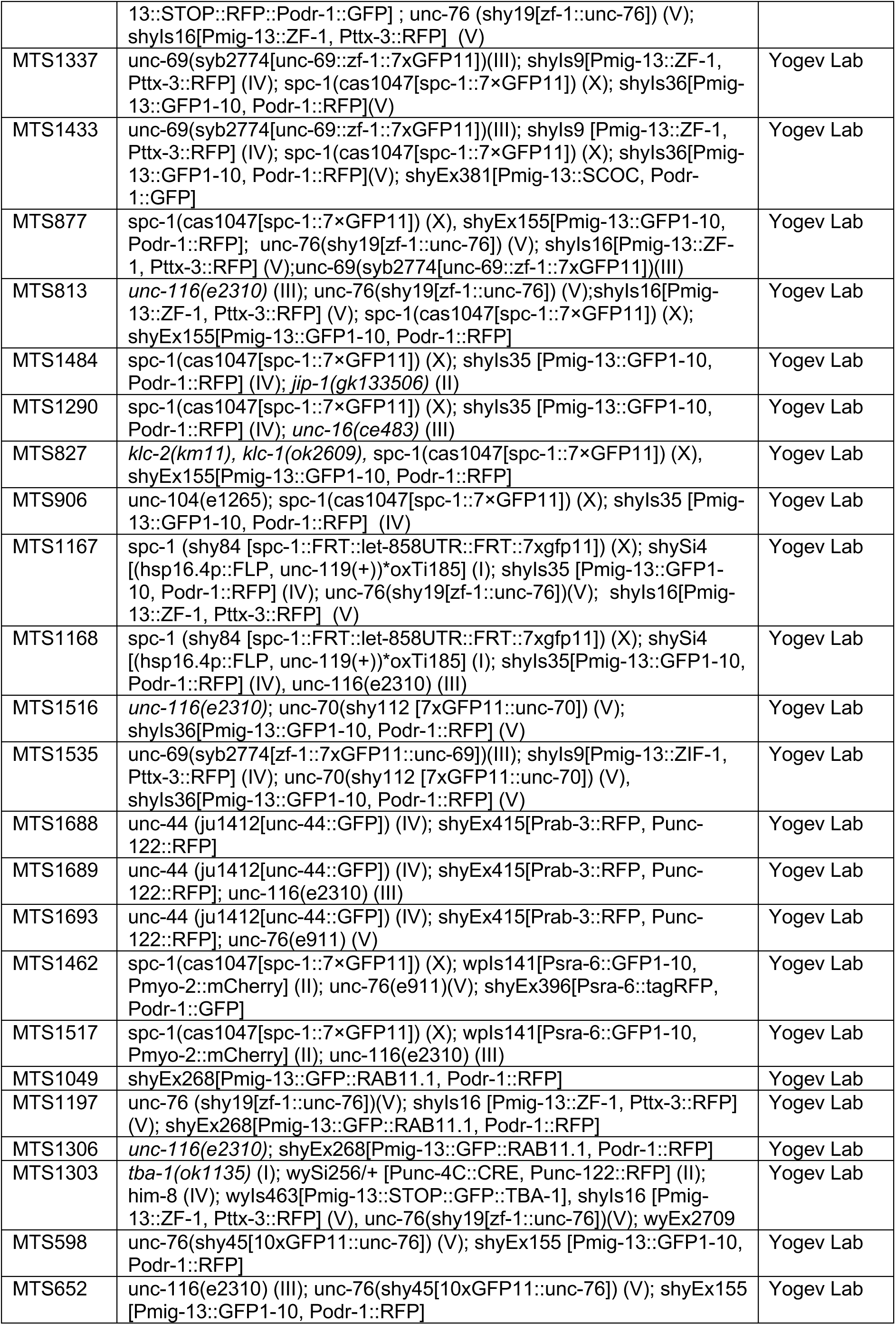
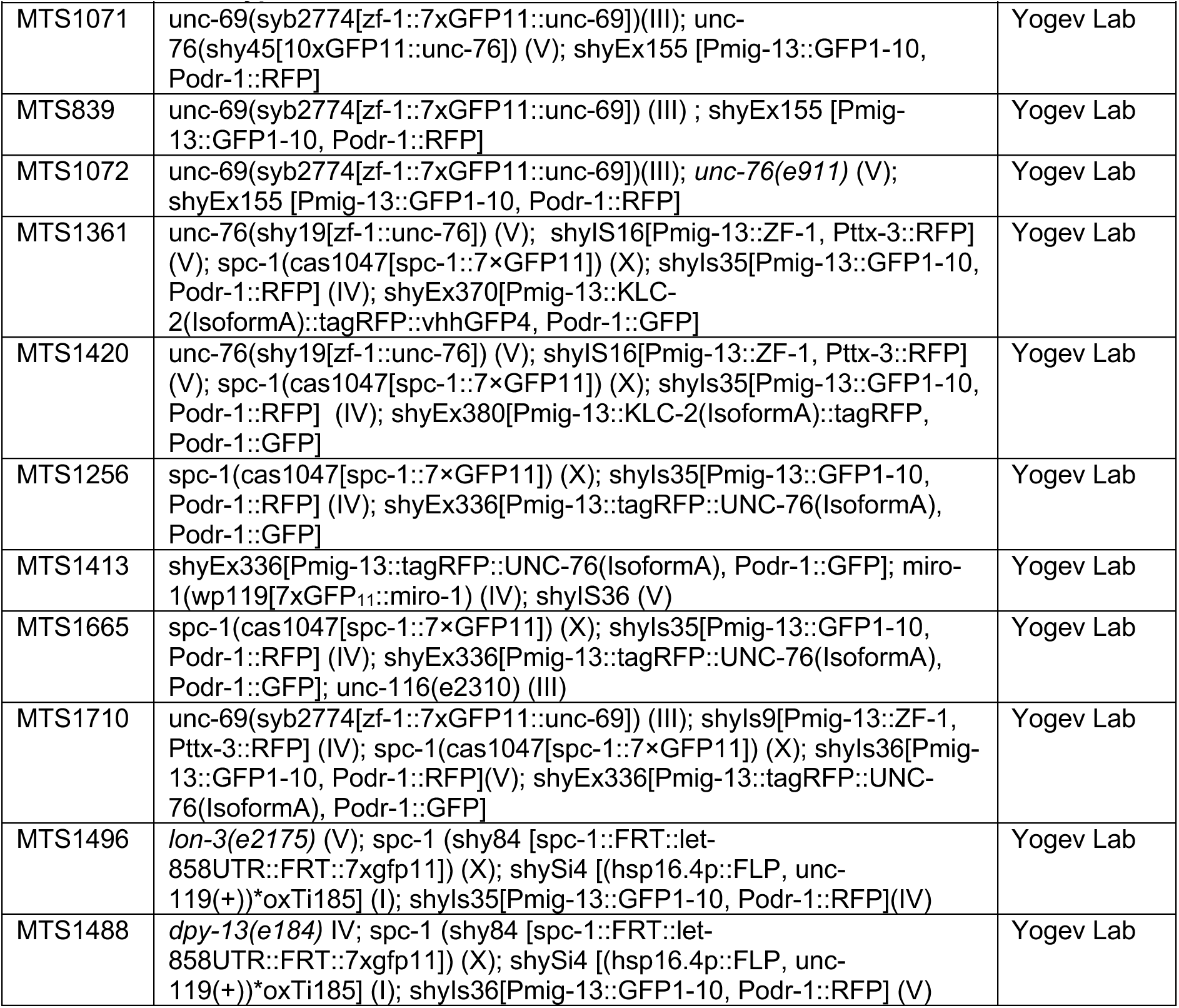

### Gene-editing by CRISPR/Cas-9

All CRISPR based gene-editing experiments were carried out based on a recently updated protocol by the Mello Lab ^28^. In brief, in a total injection mix of 20 μl, 0.5 μl Cas9 (*S. pyogenes* Cas9 3NLS, 10μg/μl, IDT# 1081058), 3.7 μg gRNA (1.85 μg/g for each RNA, if two different gRNAs were used), 900fmol dsDNA repair template or 2.2 μg of ssDNA repair template having 32-35bp homology arms and 800 ng PRF4::rol-6(su1006) plasmid was added as a selection marker ^29^. Nuclease free water was added to a total volume of 20 μl. The gRNA was synthesized from ssDNA using the EnGen sgRNA Synthesis Kit, S. pygenes (NEB# E3322S) and subsequently purifying the gRNA using a Monarch RNA cleanup kit (NEB#T2040). In case of dsDNA as a repair template, the dsDNA was melted and subsequently cooled before adding to the mixture.

The mixture was centrifuged at 14.000 rpm for 2 min and 18 μl were transferred to a fresh reaction tube and placed on ice until injected. F1 non-rollers were genotyped to find heterozygous edited mutants and singled to get homozygotes. Homozygotes were genotyped and sent for sanger sequencing.

The alleles unc-70(syb1684[unc-70(1166R + AID + 1167D)]) (V) and unc-69 (syb2774[zf-1::7xGFP11::unc-69]) (III) were generated by SunyBiotech.

Miro-1(wp119[7xGFP11::miro-1]) (IV) was also generated based on the updated protocol by the Mello Lab ^28^ but altered in the above-described procedure by adding the tracrRNA (IDT# 1072532) and crRNA separately to the injection mix instead of adding a preassembled gRNA.

For the single copy insertion by CRISPR, the injection mix contained the plasmid pOVG19 as a repair template (100 ng/μl), without a melting/cooling step before addition, as well as pMA122 (10 ng/μl) and pCFJ-90 (2 ng/μl) as selection markers. The mix was injected into the EG8078 containing a ttTi5605 landing site. The P0 generation was singled and grew for 10 days at 20°C. Successful injected animals generated non-uncoordinated moving F1- offspring, in which the uncoordinated movement was rescued by either a successful inserted single copy insertion or the formation of an extrachromosomal array. F1 progenies containing the extrachromosomal array were distinguishable from their single-copy inserted siblings by expressing a red head marker and a heat-shock inducible counter selection cassette. To exclude those extrachromosomal array containing offspring and enrich for offspring containing the successful single-copy insertion, the nematodes were heat shocked for 2h at 34°C, recovered for 4 h at 20°C and moving, non-red head containing F1’s were picked, singled and genotyped. F2 progenies that were homozygous for the single copy insertion were sent for sanger sequencing.

### sgRNAs

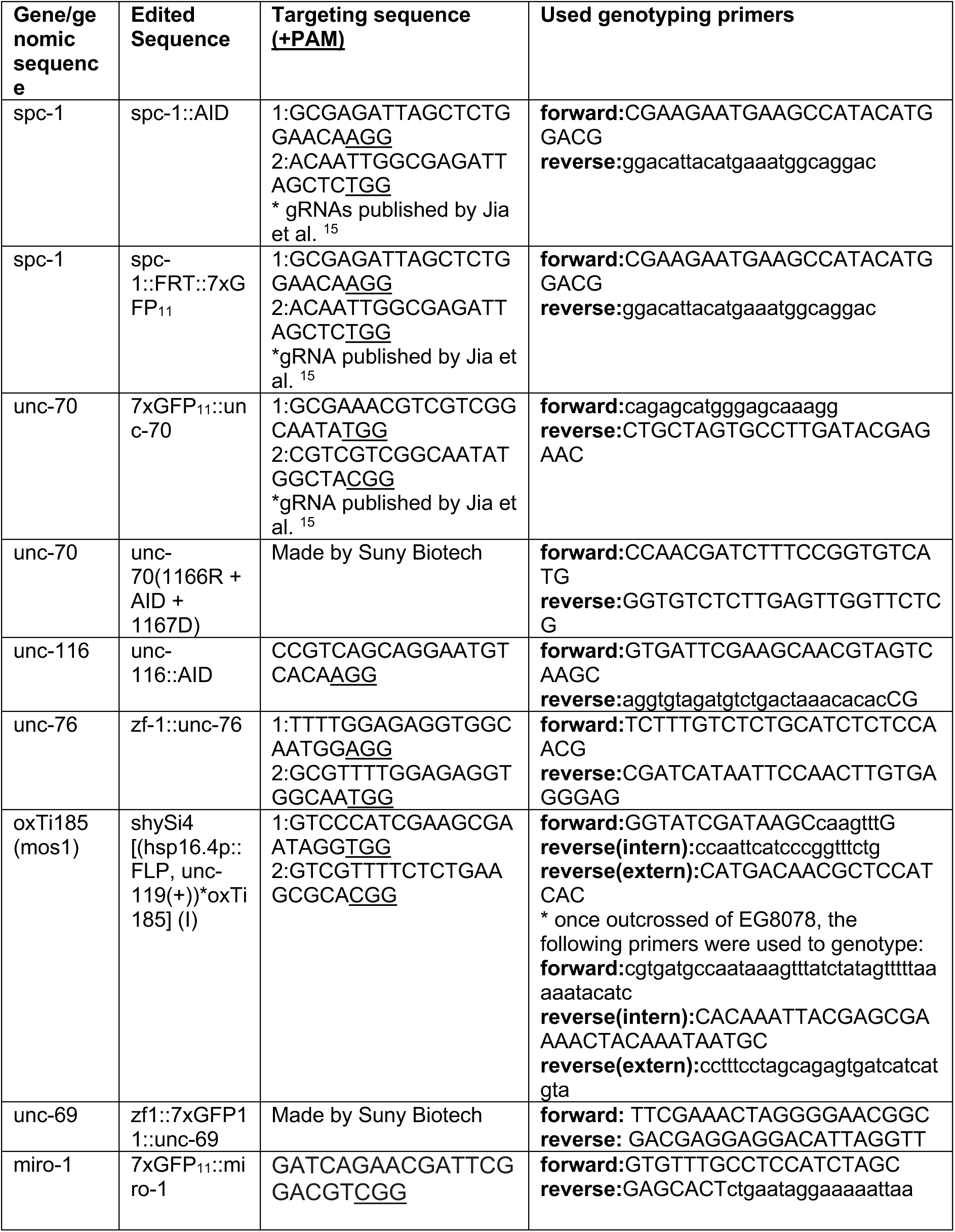

### Repair templates

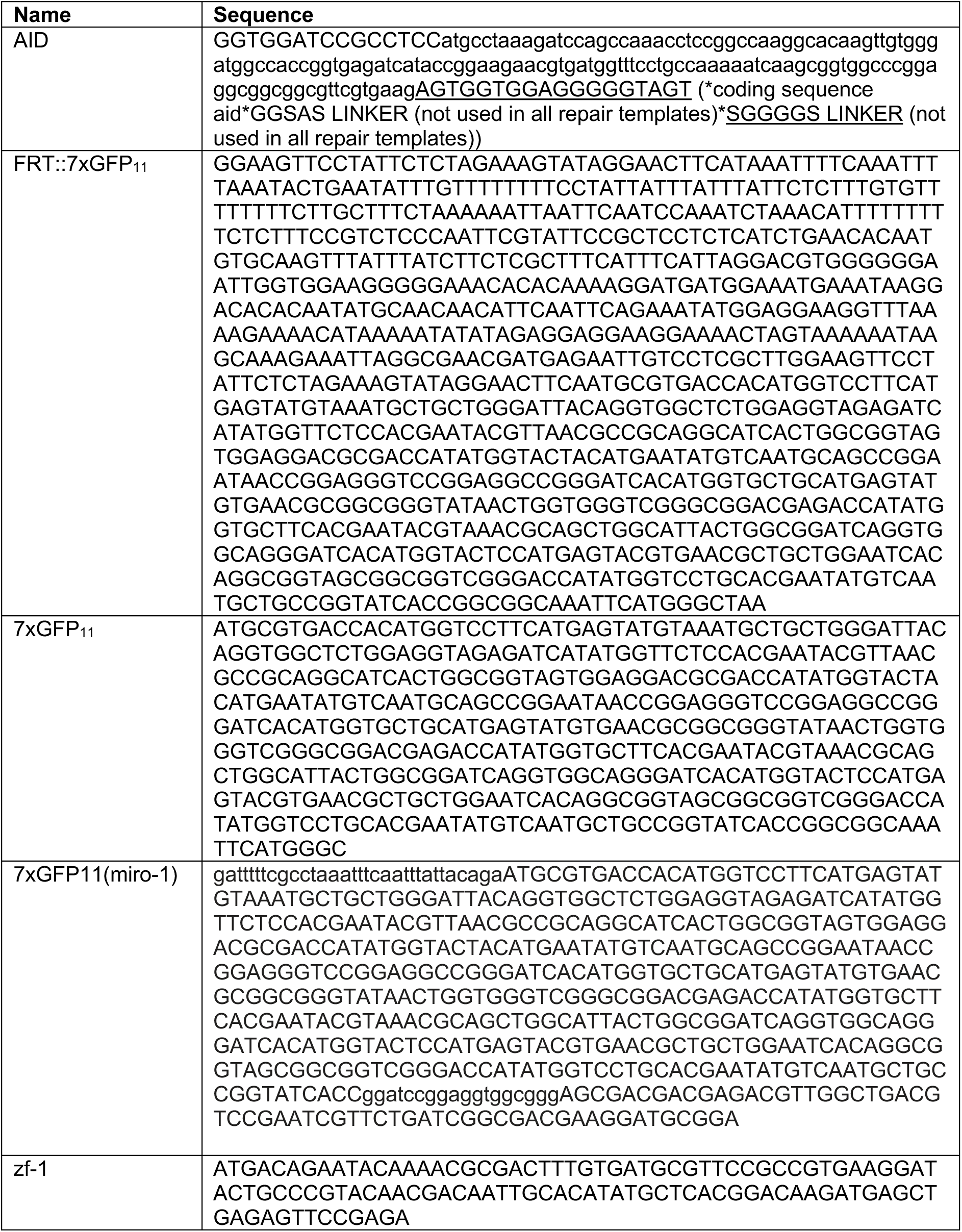

### Plasmids

Plasmids were generated by standard restriction and ligation reactions or by Gibson Assembly and confirmed by Sanger Sequencing. A list of all plasmids that were used or generated in this study is shown in the following table. Sequences and plasmids are available upon request. pGH8 was a gift from Erik Jorgensen (Addgene plasmid #19359; http://n2t.net/addgene:19359; RRID: Addgene_19359)^30^. SPTBN1 cDNA was acquired from the plasmid ß2-spectrin-HA using standard restriction and ligation procedures. ß2-spectrin-HA was a gift from Vann Bennett (Addgene plasmid #31070; http://n2t.net/addgene:31070; RRID:Addgene_31070).

### Plasmid list

| Plasmid | Injection concentration | Curator |
| --- | --- | --- |
| pRF-4[pRF4::rol-6(su1006)] | 40 ng/μl | Ambros Lab<sup>29</sup> |
| pGH8[Prab-3::mCherry::unc-54utr] | 5 ng/μl | Jorgensen Lab<sup>30</sup> |
| pOVG19[Phsp-16.4::FLP::let-858utr_pXF87 backbone] | 100 ng/μl | Yogev Lab |
| pMA122[Phsp-16.41::peel-1::tbb-2utr] | 10 ng/μl | Jorgensen Lab<sup>31</sup> |
| pCFJ90[Pmyo-2::mCherry::unc-54utr] | 2 ng/μl | Jorgensen Lab<sup>30</sup> |
| pOVG26[Pmig-13::UNC-116::SL2::mCherry] | 2.5 ng/μl | Yogev Lab |
| pOVG36[Pmig-13::tagRFP::UNC-76] | 20 ng/μl | Yogev Lab |
| pOVG43[Pmig-13::KLC-2(isoformA)::tagRFP] | 10 ng/μl | Yogev Lab |
| pOVG44[Pmig-13::KLC-2(isoformA)::tagRFP::vhhGFP4] | 10 ng/μl | Yogev Lab |
| pMR18[Pmig-13::SCOC] | 20 ng/μl | Yogev Lab |
| pMR20[Pmig-13::FEZ1] | 10 ng/μl | Yogev Lab |
| pOVG40[Pcmv::SPTBN1(Q01082-1)::eGFP] | See transfection protocols. | Yogev Lab |
| pGLS5[Pcmv::HALO::SCOC] | See transfection protocols. | Yogev Lab |
| pOVG46[Pcmv::mCherry::FEZ1] | See transfection protocols. | Yogev Lab |
| pGLS25[Pcmv::SNAP::FEZ1] | See transfection protocols. | Yogev Lab |

### Heatshock activation to temporally control fluorescence labelling of SPC-1

Heatshock was carried out at 34°C for 30 min. in a water bath incubating the nematodes in a parafilm sealed petri dish with the nematodes close to the water. The animals were subsequently placed at 20°C until imaging.

### Auxin induced single cell specific knock down

All auxin induced knock down experiments were performed by culturing the animals on plates that contain NGM supplemented with 1 mM Auxin (Alfa Aesar #A10556) and seeded with OP50.

Adult animals were allowed to lay eggs for 2h. Subsequently, adult animals were removed and the eggs were cultured at 20°C until the animals reached L4 stage and were imaged, unless indicated otherwise. The knock down condition was compared to a control condition that did not induce a knock down. As a control, i) in animals that expressed the TIR1 F-box protein from an extrachromosomal array, animals that lost the extrachromosomal array but co-cultured on auxin plates with their siblings were imaged or ii) in animals that expressed TIR1 from an integrated array, animals were cultured on NGM plates lacking auxin as a control condition.

### Microscopy

For standard microscopy, the nematodes were mounted on a 2 % agarose pad and paralyzed in a droplet of 10 mM Levamisole diluted in M9.

### Measuring animal length during development

To determine the animal length during development, adult animals were allowed to lay eggs for 30 min before removal. Removal of the adult animals set the timepoint 0 and animals were taken for imaging at 14h, 20h, 24h, 34h, 38h and 48h. Images were acquired on an upright LeicaDM4 B microscope equipped with a Leica MC170HD camera and a 10x air objective.

### Fluorescence microscopy

Animals were imaged at larval stage L4 or as 1-day old adults. To image 1-day old adults, L4 larvae were synchronized at late L4 stage and then grown for additional 12 hours. For time-lapse recordings, the animals were first incubated in a droplet of 0.5 mM Levamisole diluted in M9 for 30 min and subsequently shifted into fresh M9 medium without paralytics onto a 10% agarose pad. Images were acquired with a Laser Safe DMi8 inverted microscope (Leica) equipped with a VT-iSIM system (BioVision) and an ORCA-Flash 4.0 camera (Hamamatsu) controlled by MetaMorph Advanced Confocal Acquisition Software Package. The microscope was equipped with an HC PL APO 63x/1.40NA OIL CS2, a HC PL APO 40x/1.30NA OIL CS2, a HC PL APO 20x/0.8NA AIR and HC PL APO 100x/1.47NA OIL objective as well as 488 nm, 561 nm and 637 nm laser lines.

### Fluorescence quantification

Raw images were processed and analysed with Fiji/ImageJ v2.3.0/1.53f51^32, 33^. Images were acquired as single layer or multiple layers and then stacked into maximum projections. To measure the signal intensity along the entire axon or animal, which could not be acquired in a single field of view, multiple images were taken and stitched into a single image using the pairwise-stitching plugin with a linear blending fusion method ^34^. Time lapse microscopy images were stabilized with StackReg using the rigid body transformation method to correct for the movement or drift of animals during the recording time ^35^.

To calculate the signal intensity along a neurite, 5pxl thick lines were drawn along the length in the center of the neurite and a signal intensity profile was generated using the plot profile function.

The signal intensity was calculated as absolute signal intensities in arbitrary units as I =I_ROI_-I_Background_ or as relative intensity as I= (I_ROI1_-I_Background_)/(I_ROI2_-I_Background_) as indicated in the figures, with I_Background_ always being the background fluorescence of non-neuronal tissue next to the neuron.

### Determining the spectrin signal coverage in the axon

To calculate the portion of the axon that contained SPC-1::GFP or GFP::UNC-70, fluorescence profiles along the center midline of the axon were acquired and the fluorescence intensity along the axon was normalized to the total fluorescence intensity. This intensity ratio was binned in 5 % intervals so that each interval should contain 5 % of the total signal intensity for a perfect homogenously distributed signal. The signal was considered as absent, when the signal intensity in an interval dropped below 2% and the intervals were summed up to this drop to determine the signal coverage. The length of the signal coverage in μm was determined by multiplying the signal coverage in % with the total length of the linescan.

### Fluorescence recovery after photobleaching

Experiments were performed on a spinning disc confocal microscope (Olympus BX61) equipped with a Hamatsu C9100-50 camera and 405 nm, 488 nm and 561 nm laser lines and controlled by the software Volocity 2.0 (Quorum technologie).

The animals were prepared as just described for live cell imaging. A 2.41 μmx2.41 μm (10x10 pxl) large region of the axonal asynaptic zone was bleached using a 488nm laser, 57% laser intensity, 100 ms pulses in 60 cycles. controlled by a Perkin-Elmer photokinesis unit. 4-6 images were acquired before and 100 images were acquired after the photobleach in time intervals of 1 s of a single z-plane. To reduce acquisition bleach to a minimum, the signal intensity was binned by 2. Images were analyzed in Fiji/ImageJ and the intensity was measured by drawing a ROI i) in the region of interest, ii) in the shaft of the axon, as distant to the bleached region as possible, iii) in the background (non-neuronal tissue inside of the worm) and using the multi-measure function of Fiji. After subtracting the background signal (iii) from the intensity signal in the region of interest (i) as well as from the axon shaft (ii), a double normalization method was applied to determine the fluorescence intensity over time in the region of interest after photobleaching^36^:

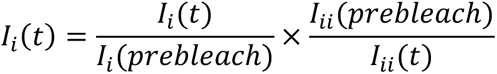

with: I_i_= signal intensity in the region of interest after background subtraction, I_ii_= signal intensity in the axonal shaft after background subtraction and t= time. Intensity signals in each region from 4-6 images before the bleach were averaged to acquire I(prebleach). Animals which moved during the image acquisition were removed.

The FRAP curves were fitted in Graphpad Prism (version 7) using a one-phase association to determine the signal intensity at which the signal plateaued. FRAP curves that could not be fitted by a one-phase association because the confidence interval of the fit was unreliable, were excluded from the analysis. The mobile phase was calculated by subtracting the signal intensity at the first timepoint after bleaching (t_0_) from the calculated plateaued signal intensity derived from a one phase association-fit of each curve (*mobile phase = I(plateau) − I(t_0_)*).

### Co-localization analysis

To quantitatively compare the co-localization between SPC-1::GFP and RFP::UNC-76 in wildtype, *unc-116(e2310)* and *unc-69 scKD* backgrounds, the ImageJ plugin “coloc 2” was used. The cell body in the green channel was encircled using the free hand selection tool and chosen as a ROI. Colocalization was scored based on the Pearson-correlation coefficient above Costes threshold to remove background signal by choosing the following parameter: Threshold regression by Costes, PSF=3 and Costes randomisations = 10.

### Analysis of spectrin peaks (hotspots)

Animals were imaged after recovering for 8 h from the heat shock. Images were taken over a time course of 1h, acquiring an image of 3 focal planes every 30 s and setting the asynaptic and synaptic zone of the DA9 neuron in the center field of view. An intensity profile was measured along the centerline of the axon at each consecutive timepoint in n=9 animals (from multiple acquisition days). The signal intensity increase of SPC-1^Flip-on^::GFP in the asynaptic-synaptic zone of the axon was determined and individual spectrin peaks were analyzed. The signal from the temporally controlled spectrin labeling probe (SPC-1^Flip-on^::GFP) at each timepoint was normalized to the signal intensity decrease over time of a constitutive labelled spectrin probe (SPC-1::GFP) acquired under the same microscopy settings of n= 3 animals to account for photobleaching during image acquisition. The loss in signal intensity of the constitutive labelled spectrin probe followed a simple linear regression model (I(t)=- 0.4362*(t)+197.3). To determine the spatial intensity along a single intensity profile over time, intensities were binned in 5 μm intervals along the axon.

To characterize individual spectrin peaks over the time course, the signal intensity profiles were exported to Matlab. The center of each individual peak along the intensity profile was refined by applying a gaussian fit. Individual peaks were identified using Matlabs find peaks algorithm. The width of each peak describes the full width at half maximum of the peak.

### Correlation in signal intensity enrichment at the distal DA9 tip

To determine the correlation between the signal enrichment of RFP::UNC-76 with either SPC-1::GFP or MIRO-1::GFP, the signal intensity at the distal DA9 tip was normalized to the signal intensity in the axonal shaft for either fluorescence channel.

The obtained relative fluorescence enrichment at the axonal tip for each channel was correlated using a spearman-correlation.

### Cell culture, transfection, and live cell imaging of COS7 cells

COS7 cells were cultured in DMEM containing 10% FBS, 1 mM sodium pyruvate, 100 U/ml penicillin, 100 mg/ml streptomycin, and 2 mM L-glutamine (all Gibco) in incubator maintained at 37°C and 5% CO_2_. 48 hours prior to imaging experiments, cells were lifted with trypsin and plated on glass-bottomed dishes (MatTek) at a concentration of 70 × 10^3^ cells per dish in DMEM without penstrep. 24 hours prior to imaging, cells were transfected with Lipofectamine 2000 (Invitrogen) in Opti-MEM (Gibco) for 2 hours. Just before imaging, Halo and SNAP tag ligands (JF549-HaloTag Ligand, JF646-SNAP tag ligand; kind gift from L. Lavis (Janelia Research Campus, Ashburn, VA)) were incubated at 1:10,000 concentration for 30 minutes. After the incubation, the imaging dish was gently washed with pre-warmed DMEM twice. Then the growth medium was replaced with live-cell imaging solution (Life Technologies). All live-cell imaging was performed at 37°C and 5% CO_2_. Spinning-disk confocal microscopy was performed using an Andor Dragonfly system equipped with a plan apochromat objective (63×, 1.4 NA, oil) and a Zyla scientific CMOS camera. Detailed protocol for live cell imaging was deposited in protocols.io^37^.

### Immunoprecipitation of SPTBN1, FEZ1, and SCOC

HEK293T cells plated on 10-cm dishes were transiently transfected with 12 μg plasmids in two separate conditions: GFP (6 μg), Fez1-Cherry (3 μg), Scoc-Halo (3 μg), and βII-Spectrin (SPTBN1)-GFP (6 μg), FEZ1-Cherry (3 μg), SCOC-Halo (3 μg). Plasmids DNA were added to 500 μl Opti-MEM and 36 μl Fugene 6 transfection mix and incubated for 20 min before adding to the cells. After 48 h of transfection, cells were rinsed twice with cold PBS, lysed in cold lysis buffer (50 mM Tris, pH 7.4, 150 mM NaCl, and 1% Triton X-100) along with protease and phosphatase inhibitors (Complete Mini-EDTA free, PhosSTOP), and centrifuged at 14,000 rpm for 8 min at 4°C to collect the supernatant, leaving behind the insoluble pellet. For the immunoprecipitation, 5000 µg protein lysates were added to 25 μL of GFP-Trap beads (Chromotek) and incubated for 1 h at 4°C on a rotating shaker. Finally, the beads were washed five times with lysis buffer and eluted in 50 μl of 2× Laemmli buffer containing 1% 2-mercaptoethanol (Sigma-Aldrich) by heating at 95°C for 5 min.

For immunoblotting, protein samples were first separated on two gels. One gel to detect GFP and spectrin-GFP, and another gel to detect Fez1 and Scoc. For immunoblot, 10 μg of proteins lysate as input and 25 μl of immunoprecipitate were loaded per gel (4–12% gradient Mini-PROTEAN TGX precast polyacrylamide gels; Bio-Rad). After electrophoresis, samples were transferred to nitrocellulose membranes using a wet blot transfer system (Bio-Rad). The membranes were blocked with 5% nonfat dry milk prepared in TBST (Tris-buffered saline containing 0.1% Tween-20) for 1 h at room temperature followed by incubation with primary antibodies diluted in 5% Milk prepared in TBST at 4°C overnight. The antibodies used were anti-GFP (goat) peroxidase conjugated, 1:10000(Rockland, #600-103-215), Anti-RFP (Rabbit), 1:3000 (Rockland, #600-401-379), anti-HaloTag (Rabbit), 1:3000 (Promega, #G9211).

The next day, the membranes were washed 5 times with TBST and the membrane with Anti-Halo and Anti-RFP was incubated with anti-Rabbit HRP-conjugated secondary antibody 1:3000 (Cell signaling, #7074S) diluted in 5% nonfat dry milk in TBST for 1 h at room temperature. After secondary antibody incubation, the membrane was washed five times with TBST. Finally, the membranes were developed using SuperSignal™ West Pico PLUS Chemiluminescent Substrate (Thermo Fisher Scientific, # 34579) and SuperSignal™ West Femto Maximum Sensitivity Substrate (Thermo Fisher Scientific, # 34094) using a Chemidoc imaging system (Bio-Rad).

### Statistical evaluation

Statistical evaluation was performed with GraphPad Prism (version 7). Each dataset was first tested for normality distribution by using the D’Agostino and Pearson test to judge the use of non- vs parametric statistical tests. Each test used in a given experiment as well as sample sizes are indicated in the figure or figure legend. Violin plots show the median as well as the 25% and 75% quartiles. All correlation analysis is based on Spearman (non-parametric) or Pearson (parametric) correlation.

